# Microaerobic conditions and the complex regulation of natural competence in *Staphylococcus aureus*

**DOI:** 10.1101/2022.04.06.487305

**Authors:** Shi Yuan Feng, Yolande Hauck, Fedy Morgene, Roza Mohammedi, Nicolas Mirouze

## Abstract

To perform genetic transformation, one of the three main Horizontal Gene Transfer mechanisms leading to virulence and antibiotic resistance genes acquisition, bacteria need to enter a physiological differentiated state called natural competence. Diverse environmental and cellular cues have been shown to trigger complex regulatory pathways ultimately activating central competence regulators controlling the expression of the genetic transformation genes. Interestingly, new bacteria displaying such aptitude are often discovered, and one of the latest is the human pathogen *Staphylococcus aureus*. However, a clear understanding of the environmental signals, regulatory pathways and central regulators involved in the development of competence in *S. aureus* is not yet available.

Here, using an original optimized protocol to naturally induce competence in planktonic cells, leading to improved transformation efficiencies (up to 5.10^−6^), we showed that the three putative central competence regulators identified in *S. aureus* are all essential for a complete development of competence. We also found that genes involved in genetic transformation can be divided in several classes depending on the regulators controlling their expression. In addition, we showed that oxygen availability is an important signal leading to competence development through the induction of one of the three central regulators. Our results demonstrate the complexity of competence development in *S. aureus*, in comparison to other historical model organisms. We anticipate our findings to be a starting point for proving the importance of natural competence and genetic transformation for *S. aureus* genomic plasticity. Furthermore, we also believe that our data will allow us to elucidate the environmental conditions leading to antibiotic resistance acquisitions *in vivo*, in this important human pathogen.

**Introductory paragraph:** *Staphylococcus aureus* has become over the years an important public health concern, due to a large range of infections and the emergence of outbreaks associated to antibiotic multi-resistant strains. *S. aureus* remarkable adaptive powers have been acquired through the acquisition of new genetic sequences, thanks to Horizontal Gene Transfer (HGT). The recent demonstration of the capacity of *S. aureus* to induce natural competence for genetic transformation prompted scientists to investigate this important mode of HGT in this new model organism in the field. Few key reports have already established that the development of competence in *S. aureus* might be complex, involving several potential master regulators, activated in response to multiple environmental signals and regulatory pathways.

In this study, we deciphered, thanks to the design of an optimized protocol, the complexity of the regulatory pathways leading to the development of competence as well as its true potential for *S. aureus*’s genomic plasticity *in vivo*. In addition, we clearly demonstrated that natural competence develops in response to oxygen limitation, a key environmental signal for a facultative anaerobe organism such as *S. aureus*.

## Introduction

A highly adaptive commensal organism such as *Staphylococcus aureus*, possesses an array of genes that allows the bacterium to grow, infect and survive in a wide variety of ecological niches. The anterior nares are generally considered as the native ecological niche of *S. aureus*, though the bacterium can be isolated from other areas of the human body including the skin, the axillae, the groin and the gastrointestinal tract ^1^. While colonization is typically not harmful, *S. aureus* may breach innate host defenses and gain access to deeper tissues, causing a variety of superficial and invasive infections ^2^. Furthermore, as a facultative anaerobe, *S. aureus* has the ability to grow and thwart the host immune system in the presence or absence of oxygen. This capacity is particularly important for *S. aureus* as its environment is known to be or become anaerobic during the course of an infection ^3,4^.

In addition to these remarkable adaptive powers, *S. aureus* also became one of the most feared pathogens in the hospital, because of the widespread emergence of antibiotic multi-resistant strains ^5,6^. Horizontal gene transfer (HGT) of antibiotic resistance genes, from other *S. aureus* strains or even from other genera, was though for years to be exclusively mediated through conjugation and transduction ^7^. However, few years ago, the demonstration that *S. aureus* is capable of becoming naturally competent for genetic transformation ^8^, changed our way to apprehend HGT in this important human pathogen.

Natural competence is a physiological adaptation that some bacterial species develop in response to various environmental signals ^9^. In response to these stimuli, bacterial cells trigger signal transduction pathways ultimately activating central competence regulators. All these steps are controlled by the so-called early competence genes. Interestingly, central competence regulators have been identified in several model organisms as transcriptional activators ^10,11^, or alternative sigma factors ^12^. Once activated, they initiate the expression of the late competence genes among which are found all the genes essential for genetic transformation.

Importantly, three potential central competence regulators have been identified in *S. aureus*: the alternative sigma factor, SigH ^13^, and two transcriptional regulators, ComK1 and ComK2 ^14^. In addition, it has been shown that *S. aureus* is able to naturally induce competence in a chemically defined medium, called CS2 ^8^. The authors detected a maximum of 1,6% of competence-inducing cells after 8 hours of growth in CS2 medium leading to transformation efficiencies that barely reached the detection limit (around 10^−10^) for a wild type strain ^8^.

In this study, our main objective was to characterize the development of competence in *S. aureus*, from the environmental stimuli sensed by the cells, the signal transduction pathways transmitting the information to the central competence regulators and their target late competence genes. To reach such goals, we first developed a new protocol to optimize the development of competence and genetic transformation in *S. aureus* planktonic cultures. Then, using various reporter strains and global transcriptomic analysis, we analyzed the role of the three central regulators during the development of competence in *S. aureus*. We particularly showed that while SigH and ComK1 are both essential for the expression of the genes involved in genetic transformation, the three regulators (SigH, ComK1 and ComK2) are all required for the full development of the competence transcriptional program. Finally, we propose that oxygen limitation, sensed by the SrrAB two-component system, activating SigH, controls the development of competence in the human pathogen *S. aureus*.

## Results

### A new protocol to naturally induce competence for genetic transformation in *S. aureus* with high efficiency

We first decided to optimize the protocol published by Morikawa and his colleagues back in 2012 ^8^ (see material and methods for details). In order to compare our results, we used the same reporter strain, expressing the *gfp* gene under the control of the *comG* late competence operon’s promoter (P*comG-gfp*) ^8^. Briefly, the reporter strain was first streaked on a BHI agarose plate. Isolated colonies are then used to inoculate a pre-culture in BHI medium. This pre-culture was then quickly stopped in exponential growth, centrifuged, washed and used to inoculate a fresh culture in CS2 medium. From this initial culture in fresh CS2, 10-fold serial dilutions were made in closed Falcon tubes. Finally, cell density (OD600nm) and the percentage of competent GFP-expressing cells were measured through flow cytometry, in every diluted culture every 30 min. Interestingly, **Fig. 1** (and **Supplementary Fig. 1**) demonstrates how our new optimized protocol was able to induce the development of competence in up to 70% of the population in a given experiment (in reality, statistical analysis evaluated this percentage at 52 ±15 % of the population, n=12). This percentage, more than 10 times higher than in the literature, was previously calculated through the detection of GFP-expressing cells by microscopy 8. Therefore, we verified that measurements performed by flow cytometry would not introduce any bias and provide similar results than microscopy (**Supplementary Fig. 2**).

**Fig. 1.**
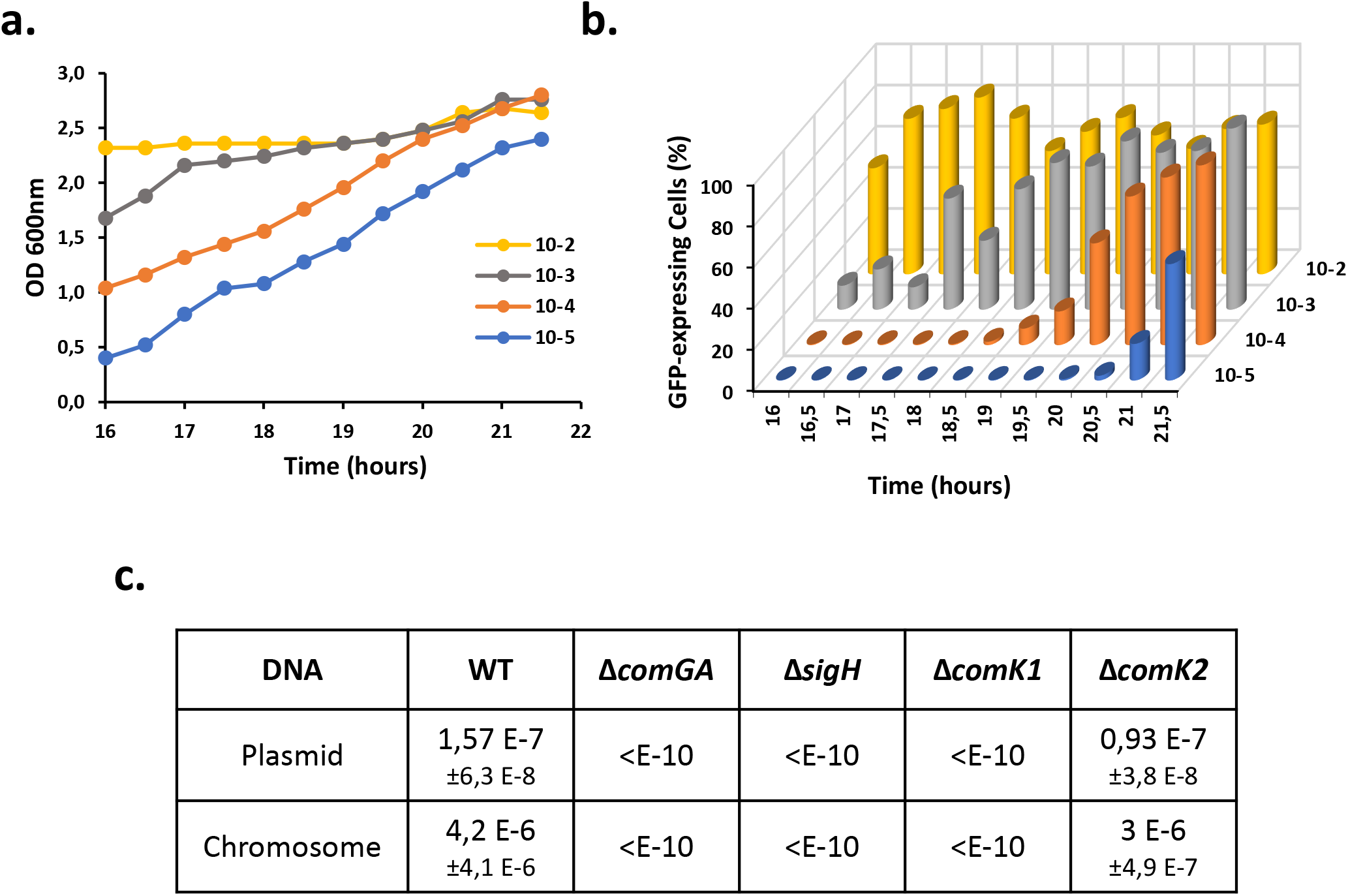
Natural competence for genetic transformation naturally develops with high efficiency in CS2 medium. a. Growth of a wild type strain expressing *gfp* under the control of the *comG* promoter (St29), in CS2 medium, between 18 and 24 hours. Dilutions from 10-2 to 10-5 are shown. The 10-2 dilution was already in stationary phase while the 10-3 dilution was entering stationary phase after 18h of growth. The 10-4 and 10-5 dilutions were respectively in late and early exponential phase at the beginning of the experiment. All the dilutions reached a similar final OD between 2,4 and 2,6. b. The percentage of competent cells was measured using flow cytometry by analyzing the percentage of cells expressing GFP under the control of the *comG* promoter. In each diluted culture, the percentage of competent GFP-expressing cells increased in late exponential phase and reached a maximum at the entry in stationary phase. c. Transformation efficiencies of wild type (N315exw/oPhi), *comGA* (St137), *sigH* (St45), *comK1* (St37) and *comK2* (St38) mutant strains using plasmidic (pCN34, KanR) or chromosomal (see material and methods for details) DNA. For each strain and each condition, the experiment has been repeated at least 10 times.

We then confirmed that our optimized protocol, associated to an improved development of competence, also led to higher genetic transformation efficiencies. Interestingly, under these laboratory conditions a wild type strain reached a transformation efficiency around 1,5.10^−7^ (±0,6.10^−^ 8) with a replicative plasmid as donor DNA **(Fig. 1c)**. Such result is 1000 times higher than what has been previously published ^8^. Importantly, the transformation efficiency even reached 4,2.10^−6^ (±4,1.10^−^ 6) when we used a *S. aureus* strain’ chromosome (harboring an antibiotic marker, see Material and Methods) as exogenous DNA **(Fig. 1c)**. To definitely confirm that the colonies growing in our assays were real transformants, we also showed that no transformation events could be detected using a strain in which *comGA*, an essential gene for genetic transformation ^15^, was deleted **(Fig. 1c)**.

### Competence naturally develops at a specific cell density and growth phase in CS2

**Fig. 1** clearly shows how competence was induced in each diluted culture using our new optimized protocol. Each additional dilution was characterized by a two-hour delay in growth. Accordingly, the development of competence was also delayed in each consecutive diluted culture **(Fig. 1** and **Supplementary Fig. 1**). However, GFP-expressing competent cells always appeared when the cultures approached OD = 2 and reached a maximum once each culture entered stationary phase.

To further verify the existence of a correlation between competence development and cell density, we repeated this experiment three times (**Supplementary Fig. 3a**). This correlation clearly showed that competence development in *S. aureus* grown in CS2 reached a maximum as the cultures entered stationary phase with an OD600nm around 2,4 (in fact between 2,2 and 2,6) (**Supplementary Fig. 3a**). High cell density, sensed by bacterial cells through quorum sensing (QS), has been proposed and verified in several model organisms as an important stress, inducing competence ^16–18^. Therefore, we then hypothesized that a QS system could be involved in the development of competence in *S. aureus*. Interestingly, two QS systems have been identified in *S. aureus* (Agr and Lux ^19^). Thus, we finally investigated the impact of *agrA* (encoding the Agr system transcriptional regulator, ^19^) and *luxS* (encoding the Lux system regulator, ^19^) genes deletion on the development of competence. Surprisingly, none of these genes were found involved in the induction of the *comG* operon’s expression (**Supplementary Fig. 3b**). This result seemed surprising, especially when compared to recent data where the deletion of *agrAC* would decrease competence by a 3-fold factor ^20^. Even though the protocols used in this study are different from ours, further investigations will be required to understand why QS does not seem involved in all conditions.

### P*comG*, P*ssb*, P*comC* and P*com*F are not controlled by the same regulators

As mentioned in the introduction, three potential competence regulators have been identified in the literature. Even though SigH has been shown to be important to activate the expression from P*comG* 8, no role has been clearly assigned to ComK1 and ComK2. Here, using our optimized protocol, we decided to test the effect of *sigH, comK1* and *comK2* deletion on the expression from four promoters: P*comG* (a promoter only activated by SigH over-expression, ^8,14^), P*ssB* (a promoter that could only be activated by ComK1 over-expression, ^14^), P*comC* and P*comF* (two promoters that could not be activated by any regulator, ^14^).

As expected, in a wild type background, the four promoters were found activated with an average percentage of GFP-expressing cells of 51,87 % for P*comG*, 77,26 % for P*ssB*, 38,32 % for P*comC* and 38,56 % for P*comF* (**Fig 2 and Supplementary Fig. 4**). When *sigH* was deleted, only the expression from P*comG* was lost (**Fig 2a**). This result confirms that SigH does not control the expression of all the genetic transformation genes and that at least one additional regulator must be involved. Interestingly, when *comK1* was inactivated, the expression from all the promoters was abolished (**Fig 2a-d**). Therefore, ComK1 is essential, alongside SigH, for the *comG* operon expression but is also absolutely required, alone, for *ssb* and *comC* expression. Interestingly, *comF* regulation seemed intermediate, as deletion of *comK1* would abolish *comF* expression while the absence of *sigH* would affect it by almost a 2-fold factor (**Fig. 3D**). Finally, in a *comk2* mutant, no effect could be detected.

**Fig. 2.**
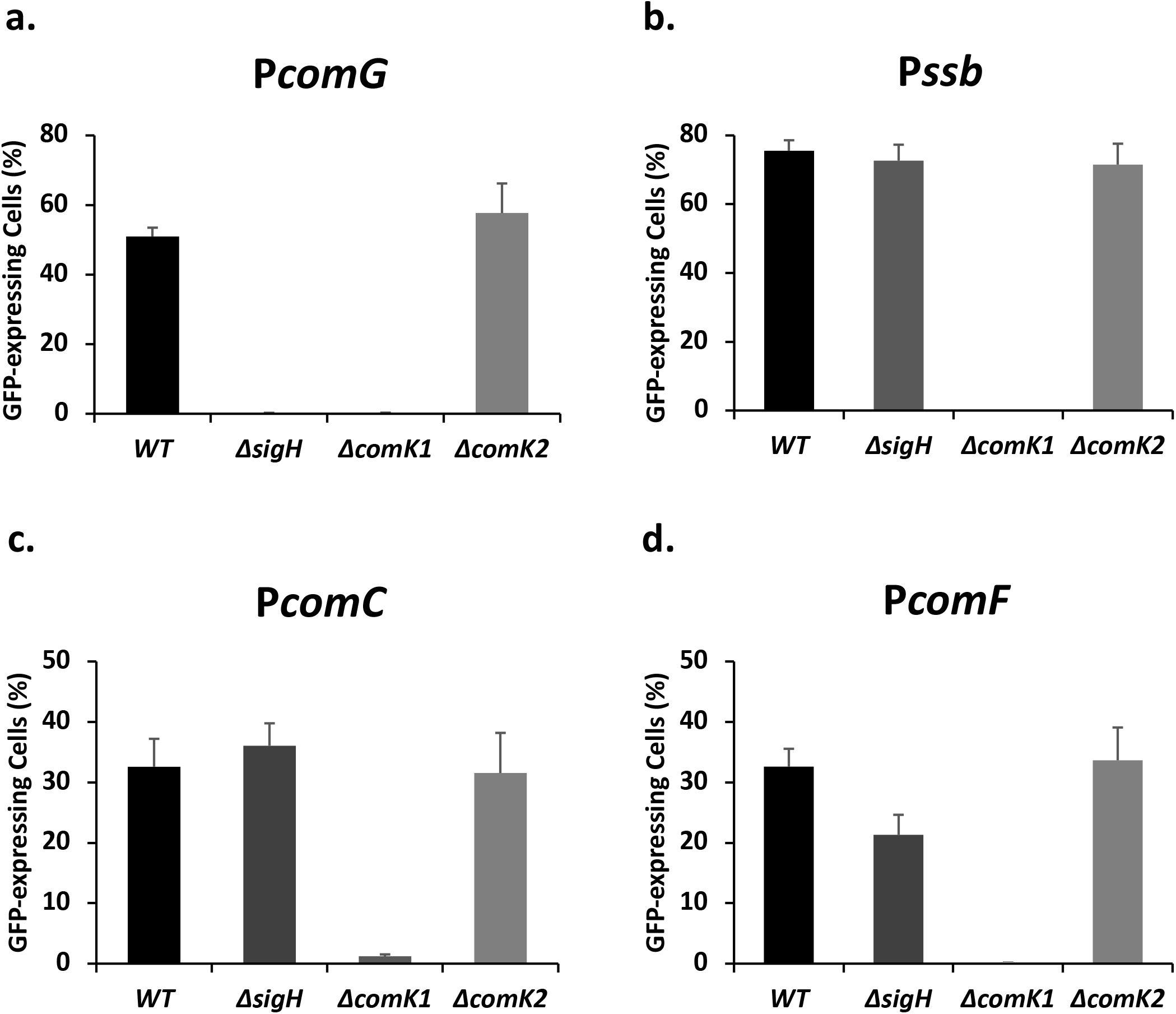
Expression of various genetic transformation-related genes is controlled by different central competence regulators. Percentage of the population expressing GFP under control of P*comG* (St29, St51, St40 and St 41) (a), P*ssb* (St50, St61, St64 and St67) (b), P*comC* (St48, St60, St63 and St66) (c) and P*comF* (St233, St235, St234 and St236) (d) in a wild type background or in the absence of *sigH, comK1*, or *comK2* was determined after 21 hours of growth in CS2 medium. Each experiment in each condition has been repeated at least 5 times.

**Fig. 3.**
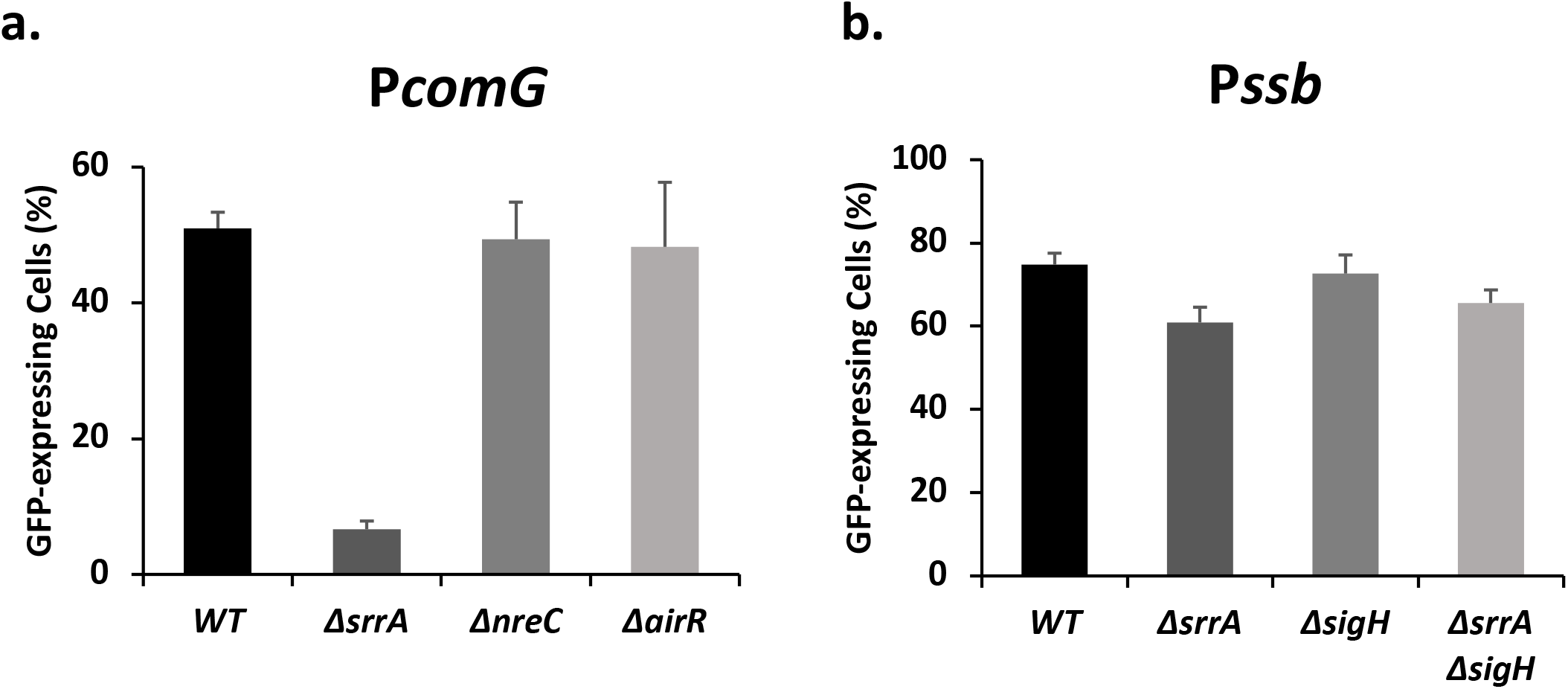
The SrrAB two-component-system controls the development of natural competence in *S. aureus*. a. Percentage of cells expressing GFP from P*comG* in wild type (St29), *ssrA* (St145), *nreC* (St158) and *airR* (St177) mutant strains. b. Percentage of cells expressing GFP from P*ssb* in wild type (St50), *srrA* (St147), *sigH* (St61) and *srrA/sigH* double mutant (St252) strains. Each experiment in each conditions has been repeated at least 5 times.

### SigH and ComK1 control natural transformation, not ComK2

To complete our results (**Fig. 2**), and exhaustively characterize the SigH, ComK1 and ComK2 regulons, we then performed a global transcriptional analysis through RNA-sequencing, by comparing the impact of the deletion of the genes encoding these three individual regulators on the competence transcriptional program.

First, we focused on the genes involved in natural genetic transformation **(Table 1** and **Supplementary Fig. 5)**. The results previously obtained were confirmed despite some small differences (discussed in **Supplementary Fig. 5**). Overall, SigH and ComK1 were both found essential for the expression of most genetic transformation genes, with the exception of *ssb*, only controlled by ComK1. Again, no role could be attributed to ComK2. Finally, we verified the transformation efficiency of strains were the genes encoding the central competence regulators were individually deleted **(Fig. 1c)**. Expectedly, only the absence of *sigH* or *comK1* abolished genetic transformation while a *comK2* mutant displayed a transformation efficiency comparable to that of a wild-type strain **(Fig. 1c)**.

**Table 1.**
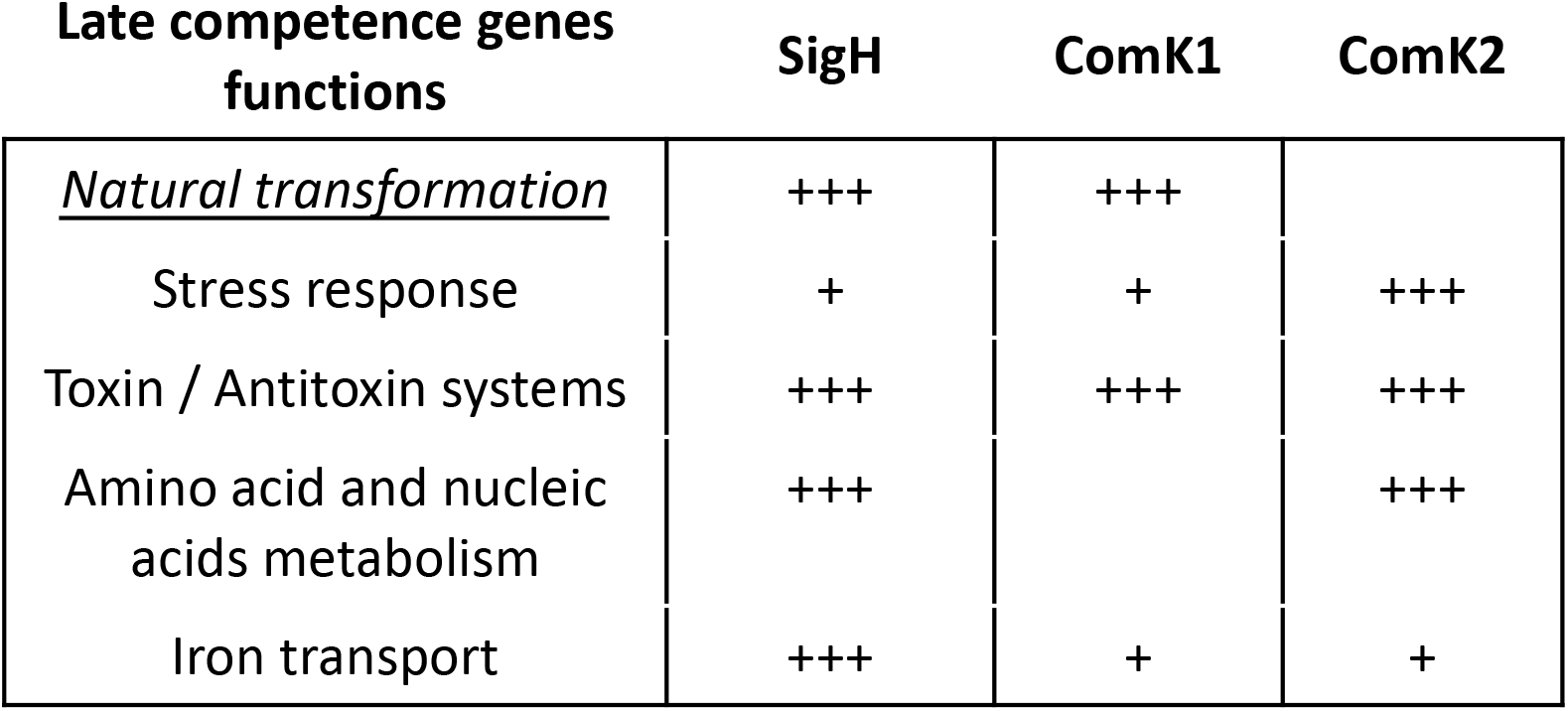
The global competence transcriptional program. Biological functions of genes induced during competence and implication of each central competence regulators. While SigH and ComK1 are both essential for the expression of the genetic transformation genes, ComK2 (alongside SigH and ComK1) is also required for a complete development of the competence transcriptional program. Indeed, in addition of genetic transformation, other cellular processes (such as the general stress response, toxin/antitoxin system, amino and nucleic acid metabolism or iron transport) are also induced during competence.

| Late competence genes<br>functions | SigH | ComK1 | ComK2 |
| --- | --- | --- | --- |
| <u>Natural transformation</u> | +++ | +++ |  |
| Stress response | + | + | +++ |
| Toxin / Antitoxin systems | +++ | +++ | +++ |
| Amino acid and nucleic<br>acids metabolism | +++ |  | +++ |
| Iron transport | +++ | + | + |

### SigH, ComK1 and ComK2 are all required for a full development of competence

In other model organisms, the competence transcriptional program always encompasses more genes than just the genes involved in genetic transformation ^21–24^. It is clearly also the case in *S. aureus*. Indeed, in comparison to the *sigH, comK1* and *comK2* mutant strains, 213/73/95 genes were respectively found overexpressed by a at least a two-fold factor in the wild type strain (**Supplementary Fig. 6**). Furthermore, if we considered higher over-expression cutoffs, the number of genes found overexpressed were still important, with 84/38/53 genes overexpressed by a three-fold factor and even 35/23/31 genes overexpressed by a five-fold factor (**Supplementary Fig. 6**). The numbers found here, especially for the three-fold expression factor, are quite similar to what has been described in other model organisms, revealing a true core set of late competence genes in *S. aureus*.

We then analyzed the functions associated to the genes induced during *S. aureus* competence transcriptional program (**Table 1, Supplementary Fig. 7, 8, and 9**). In addition to the natural genetic transformation genes, we found genes involved in (i) the regulation of the stress response (mainly but not exclusively controlled by ComK2), (ii) genes encoding multiple putative Toxin/Antitoxin systems ^25^ (controlled by the three regulators), (iii) genes involved in the amino and nucleic acids metabolism (controlled by SigH and ComK2) and (iv) genes involved in iron transport (mainly but not exclusively controlled by SigH). Importantly, our results clearly established that even though only SigH and ComK1 are essential for the expression of the genetic transformation genes, the central competence regulators (i.e. SigH, ComK1 and ComK2) are all absolutely required for the full development of the competence transcriptional program in *S. aureus*.

### Competence development is associated with virulence inhibition

Our global transcriptional analysis also revealed that the expression of numerous genes was inhibited during the natural development of competence in *S. aureus*. Indeed, we found that a total of 399 genes were inhibited in the wild type strain compared to the *sigH, comK1* or *comk2* individual mutant strains (**Supplementary Fig. 10**). This represents more than 15% of all the genes present in the N315 *S. aureus’* genome.

Interestingly, more than 10% of the repressed genes (37 to be exact), are directly involved in virulence or virulence regulation (**Supplementary Fig. 11**). Among these genes were found many well characterized virulence factors, including genes encoding for the capsule (CapA-O), serine proteases (SplA-F, SspA-C), intercellular adhesins (IcaAB), exoproteins (Hlg, Coa), surface proteins (ClfAB, fnb, FnbB, geh, sdrCD, Spa) and a lipase (Geh). In addition, the expression of an important two-component system known to drive the expression of over 20 virulence factors *in vivo, saeRS* ^26^, was found repressed during competence. Even though all the central competence regulators were found involved in virulence repression, SigH and ComK2 played a major role with an important overlap between the genes they each repress (**Supplementary Fig. 11**).

### Oxygen limitation is an important environmental signal to induce the development of competence

Finally, competence development in *S. aureus* has been demonstrated in different conditions, associated to different oxygen availability ^8^. Thus, we wondered if oxygen-sensing two-component systems (TCS) present in *S. aureus* (i.e SrrAB ^27^, NreBC ^28^ and AirRS ^29^) were involved during the early competence regulation steps in the course of planktonic growth in CS2 medium. To test such hypothesis, we compared the expression from the *comG* promoter in wild type and *srrA, nreC* and *airR* mutant strains. In this experiment, deletion of *nreC* and *airR* did not affect the expression from the *comG* promoter (**Fig. 3a**). However, when *srrA* was absent, *comG* expression was decreased by a 7-fold factor (**Fig. 3a**). Since both SigH and ComK1 are involved in the *comG* operon expression (**Fig. 2a**), we then decided to determine which central competence regulator was under the control of SrrAB. As *ssb* expression was found only controlled by ComK1 (**Fig. 2b**), we finally tested the effect of *srrA* deletion on its expression. In the absence of *srrA*, the expression of *ssb* was not affected (**Fig. 3b**). In addition, deletion of both *sigH* and *srrA* had the same impact on the *ssb* expression as the individual *sigH* and *srrA* mutant strains (**Fig. 3b**). Altogether, these results suggest that SrrAB might activate SigH to control the expression from the *comG* promoter.

Oxygen-sensing two-component systems allow *S. aureus* to survey and respond to microaerobic or anaerobic conditions. In particular, SrrAB has been shown to be a global regulator of virulence factors under low-oxygen conditions ^27^. It is therefore tempting to speculate that during *S. aureus* growth in CS2 medium, the oxygen concentration dropped in the culture. To test this hypothesis, we finally measured the concentration of dissolved oxygen throughout growth in CS2 (**Fig. 4**). Interestingly, as the OD started to increase, the oxygen concentration quickly dropped. Indeed, while the oxygen concentration stayed constant at 21% for the 8 first hours, it then decreased down to 0,27% during the next 6 hours, while the culture’s OD only reached 0,5. Few minutes after the oxygen concentration reached its lowest point, the culture reproducibly paused for roughly 1 hour, potentially to adapt to these new low oxygen conditions (**Fig. 4**). Finally, following the drop in the oxygen concentration and the pause in growth, GFP-expressing competent cells started to emerge (**Fig. 4**). Therefore, we can confidently propose that using our new optimized protocol, growth in CS2 medium is associated to a quick limitation in the oxygen availability, leading to microaerobic conditions sensed by SrrAB which in turn activates SigH for the induction of natural competence in *S. aureus*.

**Fig. 4.**
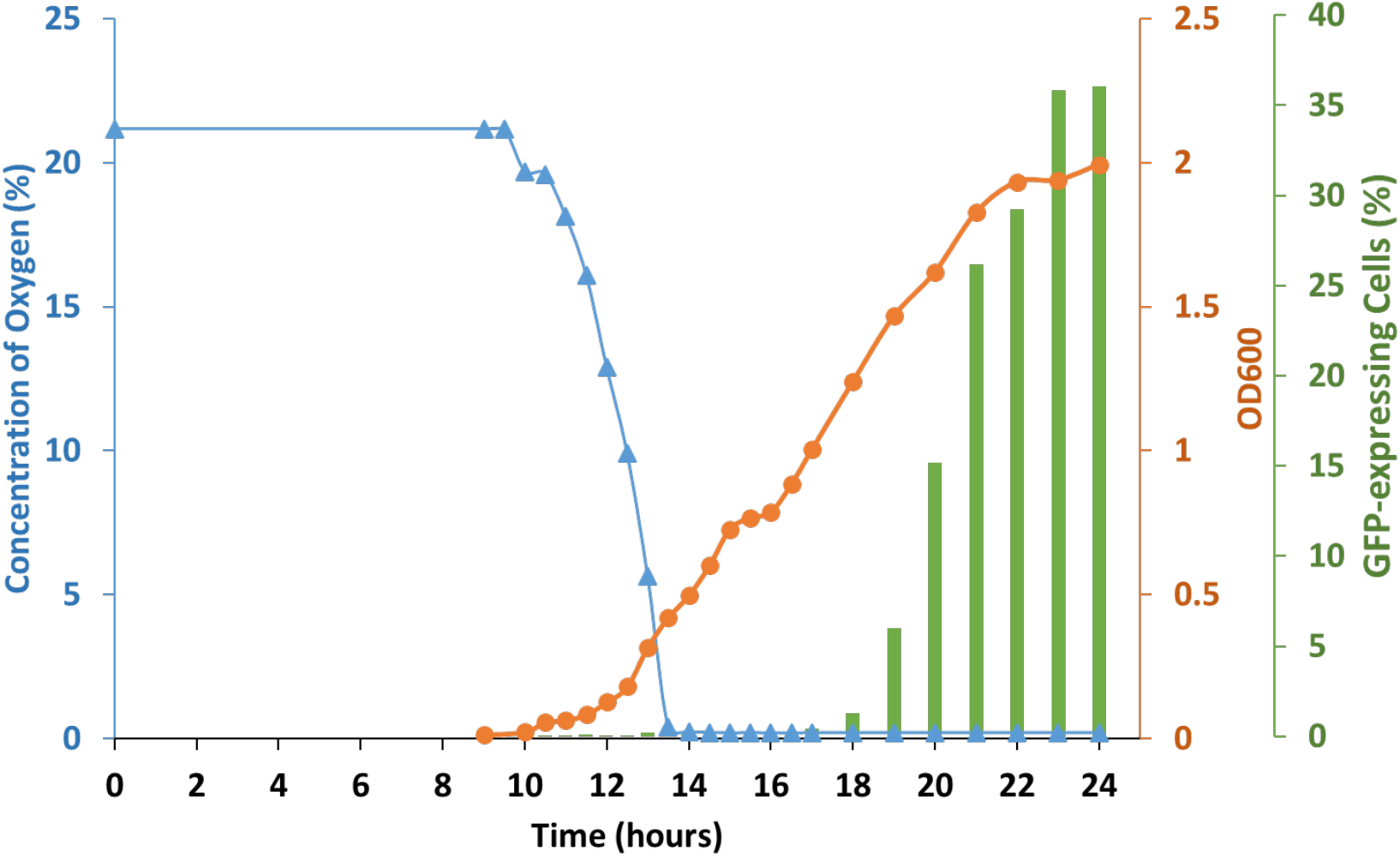
Growth in CS2 medium leads to microaerobic conditions. A wild type strain (St29) expressing GFP under the control of P*comG* was grown in CS2 medium for 24 hours. When growth became detectable, the oxygen concentration, the percentage of GFP-expressing cells and OD_600nm_ were measured every 30 min. As the OD increased, the concentration of oxygen quickly dropped from 21% to 0,27%, while the OD only reached 0,5. Importantly, our assay allows us to affirm that complete anaerobic conditions are not reached as our sensors are able to measure with confidence oxygen concentrations that are 10 times lower. Few minutes later, the growth reproducibly marked a pause, probably necessary for the cells to adapt to low oxygen conditions, which in turn induced an increase in the percentage of competent GFP-expressing cells.

## Discussion

Previously, the work of Fagerlund and her colleagues ^14^ clearly established that central competence regulators individual or combined over-expression was not enough to induce the full development of competence in *S. aureus*. In this study, we present a new optimized protocol allowing the optimum induction of natural competence in *S. aureus*. Such protocol was essential to lead the genetic study proposed here. Importantly, the resulting transformation efficiencies detected with a wild type strain, naturally inducing competence, ultimately demonstrate the true potential of this HGT mechanism to modulate *S. aureus* genetic plasticity and antibiotic resistance genes acquisition *in vivo*.

In addition, we also establish that three central competence regulators are essential for a complete development of the competence transcriptional program in *S. aureus*. While the importance of SigH was already known, we demonstrate for the first time the essentiality of ComK1 for the expression of the genes involved in genetic transformation. Importantly, we also reveal how ComK2 is involved, alongside SigH and ComK1, in the complete development of the competence transcriptional program (**Fig 5**). Indeed, our global transcriptomic study clearly shows that in addition to genetic transformation, numerous other functions are also induced during competence, a feature shared with other model organisms. It will be important to investigate in the future how these other biological processes (i.e. stress response ^30^, amino ^31^ and nucleic ^32^ acid metabolism or toxin/antitoxin systems 33,34) participate to the establishment of the competence environmental adaptation.

Furthermore, it is interesting to mention that the regulation of competence development in *S. aureus* would share traits with several important historical model organisms: *S. pneumoniae* and its alternative sigma factor (ComX), phylogenetically close from SigH ^13^, *B. subtilis* and its central competence regulator, ComK, homologous to ComK1 and ComK2 ^14^ and *V. cholerae* through the presence of several central competence regulators ^18^. Thus, it would be interesting and probably important to test in the future if other regulatory features, present in these historical model organisms, could be also used by *S. aureus* to control or modulate the development of competence.

Finally, we demonstrate how oxygen limitation, leading to microaerobic conditions, sensed by the SrrAB two-component system, controls the development of competence in *S. aureus*. Results from the literature already indicated that *S. aureus* had the ability to induce competence under different oxygen concentrations ^8,20^. Induction of competence under anaerobic conditions ^8^ or during biofilm formation 20 further reinforce this idea. We also proposed that SrrAB would activate SigH (through transcription, translation or stability) in response to variations in the environmental oxygen concentration. Additional work will be required to fully understand how a decrease in the oxygen concentration activates, directly or indirectly, SigH through SrrAB.

Importantly, microaerobic conditions are often encountered by *S. aureus in vivo*. Indeed, during an infection, energy-consuming activated neutrophils trigger oxygen deficiency while macrophages, dendritic cells and T cells induce inflammation, altering blood flow to tissues and reducing oxygen levels dramatically ^35^. In addition, oxygen-restricted microenvironments are formed during biofilm-associated infections ^36^ or abscesses ^37^. Therefore, during the course of an infection, *S. aureus* is often exposed to the environmental conditions described in this manuscript to naturally induce competence for genetic transformation, reinforcing the true potential of this HGT mode *in vivo*. However, our global transcriptomic analysis also revealed that the transcription of numerous virulence genes was repressed during the natural development of competence (**Supplementary Fig. 12 and Fig. 5**). Therefore, as an *in vivo* model, we can propose that during the course of an infection, *S. aureus* cells would induce the expression of the virulence regulon, which would ultimately lead to a local oxygen deficiency. A restricted percentage of the population would sense this environmental signal and, in response, induce natural competence for genetic transformation in a restricted part of the population. Ultimately, these competent *S. aureus* cells would repress virulence and promote HGT while the rest of the population would continue to promote the infection.

**Fig. 5.**
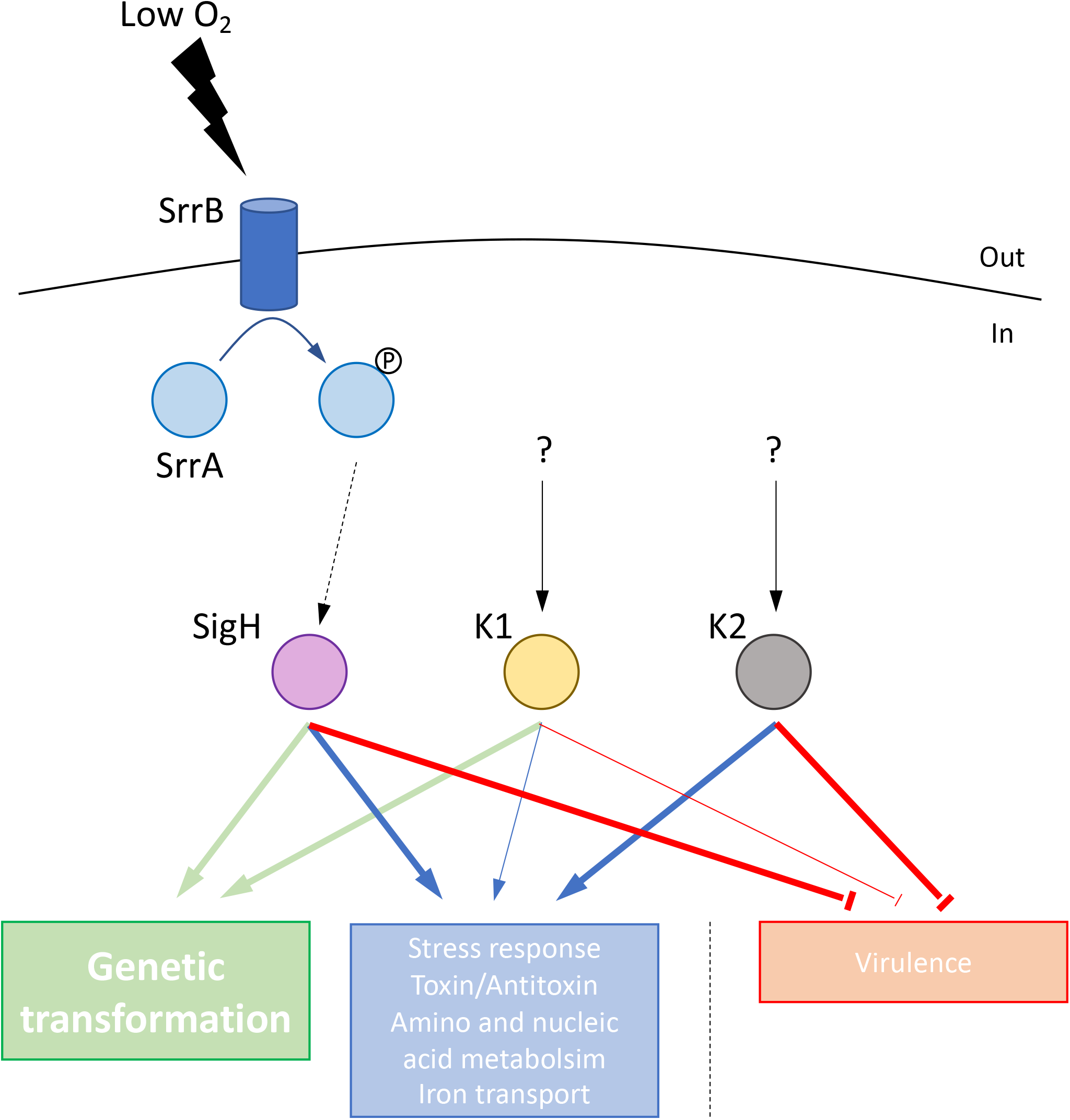
Model for the regulation of the natural development of competence in *S. aureus*. Three central competence regulators have been identified, namely SigH, ComK1 and ComK2 8,14. All three regulators are essential for a complete development of the competence transcriptional program. While SigH and ComK1 are absolutely required for the expression of genetic transformation genes, ComK2 (alongside SigH and ComK1) is also essential for the induction of additional cellular functions (i.e. stress response, toxin/antitoxin systems, amino and nucleic acid metabolism, iron transport…). In addition, the development of competence is characterized by the inhibition of virulence, mainly controlled by SigH and ComK2. Finally, we have shown that the natural development of competence is induced as the oxygen concentration in the culture is drastically decreased. This oxygen rarefaction is probably sensed by the SrrAB two-component system, which in turn activates the central competence regulator SigH.

## Materials and Methods

### Bacterial strains and culture conditions

*S. aureus* strains used in this project are all listed in **Table S1**. *Staphylococcus aureus* strains were grown in BHI medium (Becton, Dickinson and Company) or a complete synthetic medium, called CS2 ^8^, depending on the experiment. When necessary, antibiotics were used to select specific events (Kan, 200 µg/mL; Cm, 10 µg/mL).

### Optimized dilution protocol to naturally induce competence in S. aureus

Cells were isolated from -80 °C stock on BHI plate. Four clones were inoculated in 10 mL of BHI and incubated at 37 °C with shaking at 180 rpm until OD reached 2.5. This pre-culture was then centrifuged, washed in fresh CS2 medium and used to inoculate 10mL of fresh CS2 medium (OD = 0,5) in closed 50 mL Falcon tubes. From this initial CS2 culture, serial 10-fold dilutions were performed to generate 10^−1^, 10^−2^, 10^−3^, 10^−4^, 10^−5^ cultures closed 50 mL Falcon tubes, and more if needed. Several tubes for each dilution were prepared in order to take individual samples along growth (i.e. each Falcon tube was only opened once for one sample). Dilutions were finally incubated overnight at 37 °C with shaking at 120 rpm. Cells were collected when the OD was between 2 and 2.2 (GFP reporter strains, Fig. 2 and 3) or throughout growth (GFP reporter strains, Fig. 1, oxygen measurements, Fig. 4).

### Construction of GFP reporter plasmids and strains

Promoters of interest were inserted in the pRIT-GFP plasmid ^38^ using the Gibson assembly method. The promoters and the pRIT-GFP plasmid were first amplified with dedicated primers. All the oligonucleotides used in this study are listed in **Table S2**. In every primer’s tail, we designed a 25-50 bp homologous overlap regions between the extremities of each promoter and the pRIT-GFP linear plasmid. The promoters and the pRIT-GFP plasmid were amplified using the Phusion High-Fidelity DNA Polymerase (purchased from Thermo Scientific). All the PCR fragments were then purified using standard commercial silicon column (PCR cleanup kit, Macherey-Nagen) and verified by gel electrophoresis for the absence of non-specific or minor PCR fragments.

Gibson assembly master mix was prepared by adding 320 µL of 5x ISO buffer (25% w/v PEG-8000; 500 mM Tris-HCl, pH 7,5; 50 mM MgCl2; 50 mM DTT; 5 mM NAD; 1 mM each dNTP), 0.64 µL 10U/µL T5 exonuclease, 20 µL of 2U/µL Phusion polymerase, 160 µL of 40U/µL Taq DNA ligase and 699.36 µL of water (all reagents were purchased from New England Biolabs). 5 µL of the two DNA fragments mixture containing 100 ng of linear pRIT plasmid and a 3-fold excess of inserts were added to 15 µL of Gibson assembly master mix. The reaction tubes were then incubated at 50 °C for 1 hr. Finally, 1 µL of the assembly reaction was transformed into IM08B electro-competent *Escherichia coli* cells. Transformed *E. coli* cells were incubated at 37 °C on LB agar with 100 µg/mL ampicillin.

In order to select the transformants containing the expected plasmid, colony PCR were performed. The resulting plasmids were extracted from positive transformants (overnight cultures) and purified using a commercial kit (NucleoSpin Plasmid extraction kit, Macherey-Nagen). All the plasmids were verified by sequencing (GATC company). Finally, in order to obtain the final reporter strains, *S. aureus* electrocompetent cells were transformed with each constructed plasmid.

### Construction of S. aureus deletion mutants

In order to investigate the role of the main genes predicted to be involved in the regulation of competence development in *S. aureus*, listed in Table S1, allelic replacement constructs were cloned into the temperature sensitive pIMAY plasmid ^38^. All the primers used for cloning in the present study are listed in **Table S2**. Fragments corresponding to 1 kbp flanking regions of the genes to delete were amplified from *S. aureus* N315 genomic DNA using primers flanked by restriction enzyme sites compatible with the multiple cloning site (MCS) of pIMAY. The upstream or downstream regions were digested using the two chosen restriction enzymes and ligated into the pIMAY, opened with the same enzymes. The resulting plasmids were electro-transformed into the IM08B *E. coli* strain. Colony PCR was then used to verify the structure of the plasmids present in the transformants. Plasmids were purified from the positive colonies using a commercial kit (NucleoSpin Plasmid extraction kit, Macherey-Nagen). After verifying the plasmids sequence (GATC company), *S. aureus* N315ex woϕ strain was transformed by electroporation and plated on BHI agar supplemented with chloramphenicol (10 mg/mL) and incubated at 28 °C.

In order to allow pIMAY integration into the chromosome, a single colony from the transformation plate was resuspended in 200 µL of BHI. The suspension was 10-fold diluted down to 10 ^-3^ and 100 µL of each dilution was spread on BHI supplemented with chloramphenicol (10 mg/mL) and incubated at 37 °C overnight. The next day, colonies were subsequently streaked in the same conditions. Meanwhile colony PCR analysis was performed to check the absence of extrachromosomal pIMAY and whether plasmid integration had occurred in the upstream or downstream region.

Based on the colony PCR results, an overnight culture in BHI at 28 °C without chloramphenicol was performed. The overnight culture was then 10-fold diluted down to 10^−7^. 100 µL of 10^−4^ to 10^−7^ dilutions were plated onto BHI containing 1 µg/mL anhydrotetracycline (aTc). The plates were incubated at 28 °C for 2-3 days. Colonies were then patched on BHI (without antibiotic) and BHI supplemented with chloramphenicol (10 mg/mL) plates and grown at 37 °C overnight. Chloramphenicol-sensitive colonies were screened by colony PCR to identify clones containing the desired mutation. Mutants strains were finally verified by PCR and DNA sequencing.

### Flow cytometry to determine the percentage of the population expressing GFP

Following growth in CS2, 500 µL of cells were harvested by centrifugation at 11000 g for 1 min. Pellets were resuspended in 500 µL of cold 70% ethanol and incubated on ice for 20 min, in order to fix the cells. Then, *S. aureus* cells were resuspended in 500 µL of PBS (pH 7,4) after centrifugation at 11000 g for 1 min. Finally, the percentage of the population expressing GFP was evaluated by Flow cytometry (Cytoflex top-bench cytometer, Beckman-Coulter). Following Forward- and Side-scatter detection to identify individual cells, a 488 nm laser was used to distinguish GFP-expressing competent cells by comparison with the auto-fluorescence of a strain that did not express GFP (St12) (see **Supplementary Fig. 2a**).

It is important to mention that GFP is a very stable protein. Therefore, once the maximum percentage of competent cells was reached, this number stayed constant for hours. This feature does not mean that competence stays ‘open’ for hours but rather that once the maximum is reached, no new competent cells appear.

### Microscopy to determine the percentage of GFP-expressing cells

Following growth in CS2, cells were harvested and treated as explained above (see flow cytometry). Fluorescence images of cells were taken using a confocal laser-scanning microscope (ImagerieGif platform). GFP was excited at 488 nm using the blue laser and fluorescence images were collected using the green channel. Images were reconstituted using the ImageJ software.

### Natural transformation of competent S. aureus cells

Wild type strain (St12) as well as *comGA* (St137), *comK1* (St37), *comK2* (St38) and *sigH* (St45) mutant strains were first grown to competence using our optimized dilution protocol. We chose to perform the transformation experiments using the -2 dilution culture for each strain. Cells were naturally transformed following the protocol previously published ^8^ with some adjustments.

#### Transformation protocol

Briefly, at each time point (every half hour), 2 mL of cells were harvested by centrifugation at 10000xg for 1 min at 4 °C, resuspended in 2 mL of fresh CS2 and equally divided in two tubes. One or five µg of donor-DNA (plasmid or chromosome) was added to one of the tubes (the second tube is used as a “no DNA” control) and incubated at 37 °C for 2.5 hours with agitation at 180 rpm. 10 or 100 µL (tube with DNA) or 1ml (“no DNA control”) from each tube where finally mixed with 25 mL of melted BHI agar pre-cooled to 55 °C together with antibiotic, and the mixture was poured into petri dishes. After solidification, the plates were incubated at 37 °C for 48 hours. At each time point, the viability was also evaluated by serial dilution on BHI agar plates. Transformation efficiencies were finally calculated by dividing the number of transformants detected in 1 mL of culture by the total number of cells in the same volume.

Numbers presented in Fig 1C represent the mean of the highest transformation efficiencies detected along growth during each experiment. The experiments have been repeated for each strain at least 10 times to provide strong statistical relevance.

#### Donor DNA preparation

Plasmid: the pCN34 plasmid (Kan, ^39^) was used in some of the genetic transformation experiments **(Fig. 1c)**. pCN34 was purified from the St197 strain. Briefly, 50 ml of culture were harvested by centrifugation and the plasmid was purified using a plasmid purification kit (Macherey-Nagen).

Chromosomal DNA: strain St294 was used to provide donor chromosomal DNA **(Fig. 1c)**. In St294, the pIMAY-INT ^14^ plasmid (Cm) was inserted in the chromosome at the INT chromosomal site ^14^. The plasmid insertion was verified by PCR while no replicating plasmid could be detected. Briefly, 100 ml of culture were centrifuged and resuspended in 5 mL of TEG (Tris 5mM, pH8; EDTA, 10mM; Glucose, 1%) complemented with 500 µL of Proteinase K (10mg/mL), 2 mL of lysis buffer (NaOH, 0,2N; SDS, 1%) and 20 g of glass beads (Stratech, #11079-105, 0,5 mm in diameter). The cells were then broken using 5 cycles of vortex (1 min each) with 1 min in ice between each cycle. To finish cell lysis, 3mL of lysis buffer were added for 5 min at room temperature and neutralized with 6 mL of NaAc (3 M, pH 4,8). Finally, chromosomal DNA present in the supernatant was precipitated using 96% ethanol (1ml of EtOH for 500 µL of supernatant) after 2 hours of incubation at -20°C. After centrifugation, chromosomal DNA was washed using 300 µL of cold 70% ethanol. Precipitated chromosomal DNA was finally resuspended in 300 µL of Tris 5mM, pH8.

### RNA-sequencing

#### Sampling and isolation

Cultures of *S. aureus* were grown in CS2 medium, at 37°C and 180 rpm until the OD600 reached 2. To quench cellular metabolism / transcription and to stabilize RNAs, cells were harvested by centrifugation at 10000 g for 1 min at 4°C and the pellets immediately frozen in liquid nitrogen before storage at −80°C. Three independent biological replicates were collected for each of the four strains (wild type, St29; 1−*comK1*, St40; 1−*comK2*, St41; 1−*sigH*, St61). For extraction of RNA, cells were lysed using Lysing Matrix B and a FastPrep instrument (both MP Biomedicals), and RNA were isolated using the RNeasy Mini Kit (Qiagen). RNA were treated with TURBO DNase (Ambion), purified using the RNA Cleanup protocol from the RNeasy Mini Kit (Qiagen), and stored at −80°C. The integrity of the RNA was finally analyzed using an Agilent Bioanalyzer (Agilent Technologies).

#### rRNA depletion, library construction and sequencing

Removal of 23S, 16S and 5S rRNA using the RiboZero rRNA Removal Kit (Epicentre) (two times), strand-specific library construction yielding fragments of size range 100–500 bp, pooling of the 12 indexed libraries, sequencing in one flow-cell lane on a Illumina HiSeq2000 instrument with a 75 nt paired end protocol, and demultiplexing of the 12 samples of indexed reads was performed by the “Next Generation Sequencing (NGS) Core Facility” from the Institute for the Integrative Biology of the Cell (I2BC, Gif sur Yvette, France). The complete dataset from this study will be available.

#### Read mapping and analysis of differential expression

Differential expression of all annotated features was assessed using the R statistical programming environment. Differential expression was determined between the wild type strain samples (St29, n = 3) and each of the 9 samples (each n = 3) in which *sigH, comK1*, or *comK2* were absent. The output from the differential expression analysis is presented in **Supplementary Fig. 5-11**. Genes with False Discovery Rate (FDR)-corrected P-values < 0.01 and a ratio of differential expression superior to 2, 3 or 5 were considered significantly differentially expressed and are presented.

#### Oxygen concentration measurements

Oxygen concentrations were measured using the SP-PSt3-SA23-D3-OIW oxygen sensor spots (PreSens GmbH, Regensburg, Germany). These sensor spots were attached to the inner wall of 50 mL Falcon tubes with silicone glue so that the spots would always be immerged during the experiments (i.e. below the 5mL mark). The Falcon tubes were closed at T0 and remained closed for the entire experiment.

The sensor spots are covered with an oxygen-sensitive coating where molecular oxygen quenches the luminescence of an inert metal porphyrine complex immobilized in an oxygen-permeable matrix. This process guarantees a high temporal resolution and a measurement without drift or oxygen consumption. The photoluminescence lifetime of the luminophore within the sensor spot was measured using a polymer optical fiber linked to an oxygen Meter (Fibox 4 trace; PreSens GmbH). Excitation light (505 nm) was supplied by a glass fiber, which also transported the emitted fluorescence signal (600 nm) back to the oxygen meter. Briefly, an oxygen measurement was realized, through the Falcon tube plastic, by simply approaching the optical fiber from the sensor spot. At each time point, the oxygen concentration was measured three times and the results provided represent the mean of these three measurements. In our experiments, oxygen concentration was measured every 30 minutes.

## Supplementary material

### Supplementary data

Supplementary Table 1: *S. aureus* strains used in this study

Supplementary Table 2: Primers used in this study

### Supplementary Figures

**Supplementary Figure 1: Development of competence in Individual diluted cultures**.

The growth curve (in red) as well as the evolution of the percentage of competent GFP-expressing cells (black bars) is shown for the -2 (a), -3 (b), -4 (c) and -5 (d) diluted cultures of St29. Following the delay in growth associated to each dilution, the development of competence is also delayed in order to always occur as the cultures approach stationary phase.

**Supplementary Figure 2: Flow cytometry and microscopy provide identical percentages of GFP-expressing competent cells (St29)**

a. Classic flow cytometry experiment where the percentage of competent GFP-expressing cells is calculated by comparison of the fluorescence profile of a strain expressing GFP under the control of a competence-induced promoter (in this case, P*comG*, St29) and a strain that does not express GFP (left, St12). The cells showing the maximum autofluorescence in the St12 culture, provide the threshold above which cells from the St29 culture are considered as competent (marked in green).

b. Spinning-disk microscopy confirms that the percentage of competent GFP-expressing cells (St29, P*comG*-*gfp*) is increased with our optimized protocol. In St29, the competent GFP-positive frequency reached 54,1 ± 12,7% (mean ± SD, n=5). Bar = 5 μm.

**Supplementary Figure 3: Competence naturally develops at a specific cell density and growth phase in CS2**.

a. Correlation between competence development (percentage of competent cells expressing the GFP under the control of the *comG* promoter, St29) and cell density. The competence window, during which the GFP expression was at its maxima, is allowed when ODs are between 2.2 and 2.6. Data are represented as mean ± SEM of 3 independent experiments (n= 219). Each experiment is materialized by a different color (green, blue or red dots).

b. Percentage of competent cells expressing GFP under the control of the *comG* promoter in wild type (St29) or in the absence of *agrA* (St107) or *luxS* (St123) was determined after 22 hours of growth in CS2.

Each experiment in each condition has been repeated at least 5 times.

**Supplementary Figure 4: P*comG*, P*ssb*, P*comC* and P*comF* expression in all mutant strains** Percentage of competent GFP-expressing cells under the control of P*comG* (St29, St51, St40, St 41, St47, St81, St77 and St69) (a), P*ssb* (St50, St61, St64, St67, St75, St83, St79 and St71) (b), P*comC* (St48, St60, St63, St66, St74, St82, St78 and St70) (c) and P*comF* (St233, St235, St234, St236, St268, St269, St270 and St271) (d) in a wild type background or in the absence of *sigH, comK1, comK2, comK1*/*comK2, comK1*/*sigH, comK2*/*sigH* or *comK1*/comK2/*sigH* was determined after 21 hours of growth in CS2 medium. The results confirm those obtained in Fig. 2. Indeed, independently from the mutant combination considered, as long as one essential regulator is missing, the expression is inhibited.

Each experiment in each condition has been repeated at least 5 times.

**Supplementary Figure 5: Genetic transformation genes expression is induced during natural and controlled by the competence central regulators**.

Overall, we could define three classes of genetic transformation genes: (i) Class I: genes for which SigH and ComK1 are both essential, (ii) Class II: genes (*ssb*) exclusively controlled by ComK1 (and (iii) Class III genes are controlled by SigH and ComK1 even though their impact is not as important. However, compared to the GFP reporter strains (Fig 2a-d), the impact of the regulators deletion does not always provide the same result by RNA-sequencing. Indeed, the results are confirmed for the *comG, ssb* or *comF* expression, controlled either by SigH and ComK1 or only by ComK1. However, if we consider the *comC* expression, ComK1 and SigH seem to have a small impact through RNA-sequencing (Table 2), while only ComK1 appeared essential using GFP-reporter strains (Fig 3C).

Importantly, the differences observed between GFP and RNA-sequencing experiments could be explained by the fact that GFP accumulates during time (because of its stability) while mRNA contents evolve rapidly (because of a high turnover 29). Therefore, samples of GFP reporter strains could be taken later into stationary phase, while we extracted the mRNA content at a specific time during the transition to stationary phase. Interestingly, these results suggest that all the genetic transformation genes are not expressed exactly at the same time and that specific regulations (involving one or several regulators) could be at play.

**Supplementary Figure 6: Global transcriptomic analysis (RNA-sequencing) of competence development in *S. aureus***.

Each colored circle represents a regulon, for which the expression is increased at least a twofold factor, and controlled by SigH (in blue), ComK1 (in yellow), by ComK2 (in green) or by the three (outside black circle). The number of genes in each category is shown inside the circles or at the intersection between circles, when the same gene is controlled by more than one regulator. The tables outside show the number of genes for which the expression is increased by a two-fold, three-fold or five-fold factor and controlled by each central regulator (the table color corresponds to the color of the circles) or by the three regulators (in black).

**Supplementary Figure 7: SigH regulon during competence**

List of genes, and their function, for which the expression is induced by more than a three fold factor by SigH. In the last column is provided the exact expression-induction factor associated to each gene under the control of SigH.

**Supplementary Figure 8: ComK1 regulon during competence**

List of genes, and their function, for which the expression is induced by more than a three fold factor by ComK1. In the last column is provided the exact expression-induction factor associated to each gene under the control of ComK1.

**Supplementary Figure 9: ComK2 regulon during competence**

List of genes, and their function, for which the expression is induced by more than a three fold factor by ComK2. In the last column is provided the exact expression-induction factor associated to each gene under the control of ComK2.

**Supplementary Figure 10: Genes inhibited during competence**

Each colored circle represents the regulon, for which the expression is decreased at least twofold, controlled by SigH (in blue), ComK1 (in yellow), by ComK2 (in green) or by the three (outside black circle). The number of genes in each category is shown inside the circle or at the intersection between circles, when the same gene is controlled by more than one regulator. The total number of genes inhibited by SigH, ComK1 and ComK2 during competence development is shown below.

**Supplementary Figure 11: Expression of virulence-related genes is inhibited during competence**.

List of genes involved in virulence, and their function, that were found inhibited during the development of competence. The “inhibition” ratio calculated by comparison of the expression in wild type and mutant strains is presented in the last three columns. Ratios superior to 3 are in bold. ‘K1’ and ‘K2’ refer to ComK1 and ComK2.

## Acknowledgements

We thank Kazuya Morikawa and Tarek Msadek for providing *S. aureus* strains and constructs and for the scientific discussions. We also want to thank the Imagerie-Gif flow cytometry facility (Institute for the Integrative Biology of the Cell, I2BC, Gif sur Yvette, FRANCE) for their help and support.

This work was supported by a “Young Researcher grant” from the French National Research Agency to Nicolas Mirouze (ANR-18-CE35-0004 GenTranSa).

