## Supplemental data for "Microaerobic conditions and the complex regulation of natural competence in *Staphylococcus aureus*"

**Supplementary data**

Table S1. Strains

| Strains | Genotype/Construction | Source |
|---|---|---|
| St012 | N315ex woϕ | 8 |
| St029 | N315ex woϕ / pRIT-PcomG-gfp | 8 |
| St048 | N315ex woϕ / pRIT-PcomC-gfp | This study |
| St050 | *N315ex woϕ / pRIT-Pssb-gfp* | This study |
| St233 | N315ex woϕ / pRIT-P*comF-gfp* | This study |
| St037 | *N315ex woϕ ΔcomK1* | This study |
| St040 | *ΔcomK1 / pRIT-PcomG-gfp* | This study |
| St063 | *ΔcomK1 / pRIT-PcomC-gfp* | This study |
| St064 | *ΔcomK1 / pRIT-Pssb-gfp* | This study |
| St234 | *ΔcomK1 / pRIT-PcomF-gfp* | This study |
| St038 | *N315ex woϕ ΔcomK2* | This study |
| St041 | *ΔcomK2 / pRIT-PcomG-gfp* | This study |
| St066 | *ΔcomK2 / pRIT-PcomC-gfp* | This study |
| St067 | *ΔcomK2 / pRIT-Pssb-gfp* | This study |
| St236 | *ΔcomK2 / pRIT-PcomF-gfp* | This study |
| St045 | *N315ex woϕ ΔsigH* | This study |
| St051 | *ΔsigH / pRIT-PcomG-gfp* | This study |
| St060 | *ΔsigH / pRIT-PcomC-gfp* | This study |
| St061 | *ΔsigH / pRIT-Pssb-gfp* | This study |
| St235 | *ΔsigH / pRIT-PcomF-gfp* | This study |
| St044 | *N315ex woϕ ΔcomK1+ΔcomK2* | This study |
| St047 | *ΔcomK1+ΔcomK2 / pRIT-PcomG-gfp* | This study |
| St074 | *ΔcomK1+ΔcomK2 / pRIT-PcomC-gfp* | This study |
| St075 | *ΔcomK1+ΔcomK2 / pRIT-Pssb-gfp* | This study |
| St268 | *ΔcomK1+ΔcomK2 / pRIT-PcomF-GFP* | This study |
| St055 | *N315ex woϕ ΔcomK1+ΔsigH* | This study |
| St081 | *ΔcomK1+ΔsigH / pRIT-PcomG-gfp* | This study |
| St082 | *ΔcomK1+ΔsigH / pRIT-PcomC-gfp* | This study |
| St083 | *ΔcomK1+ΔsigH / pRIT-Pssb-gfp* | This study |
| St269 | *ΔcomK1+ΔsigH / pRIT-PcomF-GFP* | This study |
| St053 | *N315ex woϕ ΔcomK2+ΔsigH* | This study |
| St077 | *ΔcomK2+ΔsigH / pRIT-PcomG-gfp* | This study |
| St078 | *ΔcomK2+ΔsigH / pRIT-PcomC-gfp* | This study |
| St079 | *ΔcomK2+ΔsigH / pRIT-Pssb-gfp* | This study |
| St270 | *ΔcomK2+ΔsigH / pRIT-PcomF-GFP* | This study |
| St054 | *N315ex woϕ ΔcomK1+ΔcomK2+ΔsigH* | This study |

| | | |
|---|---|---|
| St069 | *ΔcomK1+ΔcomK2+ΔsigH / pRIT-PcomG-gfp* | This study |
| St070 | *ΔcomK1+ΔcomK2+ΔsigH / pRIT-PcomC-gfp* | This study |
| St071 | *ΔcomK1+ΔcomK2+ΔsigH / pRIT-Pssb-gfp* | This study |
| St271 | *ΔcomK1+ΔcomK2+ΔsigH / pRIT-PcomF-GFP* | This study |
| St103 | *N315ex woϕ ΔagrA* | This study |
| St107 | *ΔagrA / pRIT-PcomG-GFP* | This study |
| St122 | *N315ex woϕ ΔluxS* | This study |
| St123 | *ΔluxS / pRIT-PcomG-GFP* | This study |
| St117 | *N315ex woϕ ΔsrrA* | This study |
| St145 | *ΔsrrA / pRIT-PcomG-gfp* | This study |
| St147 | *ΔsrrA / pRIT-Pssb-gfp* | This study |
| St118 | *N315ex woϕ ΔnreC* | This study |
| St158 | *ΔnreC / pRIT-PcomG-gfp* | This study |
| St142 | *N315ex woϕ ΔairR* | This study |
| St177 | *ΔairR / pRIT-PcomG-gfp* | This study |
| St250 | *N315ex woϕ ΔsrrA+ΔsigH* | This study |
| St197 | *N315ex woϕ* pCN34 | This study |
| St252 | *ΔsrrA+ΔsigH / pRIT-Pssb-gfp* | This study |

Table S2. Primers

| Name | Nucleotide sequence | Descriptions |
|---|---|---|
| IM151<br>IM152 | TAC ATG TCA AGA ATA AAC TGC CAA AGC<br>AAT ACC TGT GAC GGA AGA TCA CTT CG | Used for verifying and sequencing the allelic replacement pIMAY constructs |
| KpnI-sigH KO-F-F<br>SalI-sigH KO-F-R | GCA GGT ACC GAC CCG CAT AAC TTG GGA TCA ATT TTA AGA<br>GCA GTC GAC CCC CTT CTA TCT AAA ATT TAA GGT TAG TTT AAT ATT GTT ACA TTC | To amplify upstream of *sigH* used for the deletion of *sigH* |
| EcoRI-sigH KO-R-F<br>XmaI-sigH KO-R-R | GCA GAA TTC AGC GCC TTA GGA CGT GAA TTG AAT TAT AAC GTG<br>GCA CCC GGG CTG GTG TTT CTC GGC CAA ACA TAT CTA CTA ATA C | To amplify downstream of *sigH* used for the deletion of *sigH* |
| sigH-OUT-F<br>sigH-OUT-R | TGC AGC GCT TAT TGC ACC ATA TGA ATA TGC<br>TTG CCC TCC CAC TCT TAA CAT TTC GTG A | To verify the *sigH* deletion mutant |
| Kpn I-SA2017-up-F<br>Sal I-SA2017-up-R | GCA GGT ACC CAT ATG ACA CTC CCA ATG C<br>GCA GTC GAC GTA GAA GTA CCT CCA AAA ATC AAT | To amplify upstream of *sa2107* used for the deletion of *sa2017* |
| EcoRI-SA21017-down-F<br>Sac I-SA2107-down-R | GCA GAA TTC GCC CAT TAA CCT ATT TTT CAT A<br>GCA GAG CTC ACA ATA TTT GAT GCC TGT GCT A | To amplify downstream of *sa2107* used for the deletion of *sa2017* |
| check-mutant-SA2107-F<br>check-mutant-SA2107-R | TGA AAG TCA GTC GTA CTC GAC AT<br>GTT GCT CCC ATA TGC ATC TCA | To verify the *sa2107* deletion mutant |
| Kpn I-K1-up-F<br>Sal I-K1-up-R | GGT ACC GAA TGT TTA GTC ATG GTA CGT TGA TG<br>GCA GTC GAC AAG CAA AAC CTC GCT TTA TTA AGT TT | To amplify upstream of *comK1* used for the deletion of *comK1* |
| Sma I-K1-down-F<br>Sac I-K1-down-R | GCA CCC GGG TAT GCG GAA TGT TTT ATA<br>GCA GAG CTC AAG GTT TGA TGT TAT CAG TGA ATG | To amplify downstream of *comK1* used for the deletion of *comK1* |
| Kpn I-K2-up-F<br>Sal I-RK2-up- | GCA GGT ACC ATC TAT GCA TTG TTA CAT ATT CAT<br>GCA GTC GAC ATA TAG TAC TCC TCG TAT AAT AAG | To amplify upstream of *comK2* used for the deletion of *comK2* |
| EcoR I-K2-down-F<br>Sac I-K2-down-R | GCA GAA TTC GAA GCA CTT CAT TGA AAA TAC<br>GCA GAG CTC AAG TGT CAG ATT CGT GTA AT | To amplify downstream of *comK2* used for the deletion of *comK2* |
| check-mutant-comK1-F<br>check-mutant-comK1-R | TCT CAG GGA ATG CTA TGG TTA AAG T<br>CAG TGG CTT ATT GGT CAC AGG | To verify the *comK1* deletion mutant |
| check-mutant-comK2-F<br>check-mutant-comK2-R | GAG ACA ACA GCA AAT ATG ACA ACA AG<br>GTG CAT GAA TAT TAC CAC TTC TAA TCG | To verify the *comK2* deletion mutant |
| Kpn I-up-agrA-F<br>Sal I-up-agrA-R | GCA GGTACC TTA GTG ACC ATG ATC ATA ATG TAT TTG AGT<br>GCA GTCGAC TTC ACA TCC TTA TGG CTA GTT GTT AA | To amplify upstream of *agrA* used for the deletion of *agrA* |
| EcoR I-down-agrA-F<br>Sac I-down-agrA-R | GCA GAATTC AAT AAG ATA ATA AAG TCA GTT AAC GGC GTA<br>GCA GAGCTC TCT GCT GAT ATG TTA TTT GAA CCA AG | To amplify downstream of *agrA* used for the deletion of *agrA* |
| Check-mutant-agrA-F<br>Check-mutant-agrA-R | GTA TTA CTT CTA TGG AAG TAG AGC CGT ATT<br>GTG TAT CGC ACG AAT GAA GCA | To verify the *agrA* deletion mutant |
| Kpn I-up srrA-F<br>EcoR I-up srrA-R | GCA GGT ACC GAC ATT GAT TGC CAA AGG ACT AGT TGA G<br>GCA GAA TTC TAC CTC CCA CAC ATG CTT TTC TTT ACA | To amplify upstream of *srrA* used for the deletion of *srrA* |
| sma I-down srrA-F<br>Sac I-down srrA-R | GCA CCC GGG GGG CGT TGG GTA TAA ATT TGA GGT TAA ATC TA<br>GCA GAG CTC GCT CAT TAG TCA TAT CAC GAA CTG TCA C | To amplify downstream of *srrA* used for the deletion of *srrA* |
| Check-mutant-srrA-F<br>Check-mutant-srrA-R | GAA ATT ATC ACA AGC AGC AAT GGA AGT ACT<br>TGG TAT TGT TAC AGA ACC GGA TGA AAT AAA AG | To verify the *srrA* deletion mutant |
| Sal I-up nreC-F<br>Sma I-up nreC-R | GCA GTC GAC GAG CAT TCA ATC GAA AAG ATA GTT TTT GCT GAT G<br>GCA CCC GGG TCA AAT TGG AAT GTT CAA TGT AAC ATT GGT AC | To amplify upstream of *nreC* used for the deletion of *nreC* |
| Not I-down nreC-F | GCA GCG GCC GC CGA GAG GAA ATG ATT ATC TTC TGG C | |

| | | |
|---|---|---|
| Sac I-down nreC-R | GCA GAG CTC GGTGACTGAATTTTGGCATACAC | To amplify downstream of *nreC* used for the deletion of *nreC* |
| Check-mutant-nreC-F<br>Check-mutant-nreC-R | GAT GAA TTA GGG TGT AAG TCA TGA TTA ATG AGG AC<br>CTATTGAAGTTGCTACAACACTTCCAGCAC | To verify the *nreC* deletion mutant |
| Sal I-up airR-F<br>EcoR I-up airR-R | GCA GTC GAC CAA GGT GCG TTA AAA TAT TTA ATT GAG GGC<br>GCA GAA TTC CAT GGG TTA TCT CCT TAA ATC AAG CTA TT | To amplify upstream of *airR* used for the deletion of *airR* |
| sma I-down airR-F<br>Sac I-down airR-R | GCA CCC GGG GTG TAA GTT TTA TCA AAT CTA GTG ATT GTG<br>GCA GAG CTC GGT GAA ACA GCA TTC TAA TTG CAA TTA | To amplify downstream of *airR* used for the deletion of *airR* |
| Check-mutant-airR-F<br>Check-mutant-airR-R | GGA GCG ATT TGA TGG AAC AAA GG<br>GGC CAA TTT TAG TTG CAA TTT CTT CGC | To verify the *airR* deletion mutant |
| KpnI-up-luxS-F<br>Sal I-up-luxS-R | GCA GGTACC AAT ATG TTG TCG TAG TTG TGC TAC<br>GCA GTCGAC TTT GAA TTT CCT CCT ATT TAC TAC TCA A | To amplify upstream of *luxS* used for the deletion of *luxS* |
| Sma I-down-luxS-F<br>Sac I-down-luxS-R | GCA CCCGGG ATC TTA GTC AAT CAA GTT AAT CAG AAA AGC<br>GCA GAGCTC AAA GCT GAA ACG CTT GAA GAA GC | To amplify downstream of *luxS* used for the deletion of *luxS* |
| Check mutant-luxS-F<br>Check mutant-luxS-R | GCC ATC TAA AAT CAC ACC ATG CAC<br>CAT TAG GTT CTC AAA TGG TTG TAC TTG | To verify the *luxS* deletion mutant |
| KpnI-up-comGA-F<br>Sal I-up-comGA-R | GCA GGTACC CGG AAG GTC CGA GTG TTA GTA AAT<br>GCA GTCGAC CAC CTC CTA CAT ATA ATC ACG TAG GAG | To amplify upstream of *comGA* used for the deletion of *comGA* |
| Sma I-down-comGA-F<br>Sac I-down-comGA-R | GCA CCCGGG ACT ACA TTC TAA GAA GCG ACA ATT AAG TAA<br>GCA GAGCTC CAT CTC TAT CAA TGT AAA CGC TTG AG | To amplify downstream of *comGA* used for the deletion of *comGA* |
| Check mutant-comGA-F<br>Check mutant-comGA-R | CTA TTG TTG ATG TGG TTG TTA TTC CAG<br>CAA CCT GTT GAT TGT ATG TGA GCA G | To verify the *comGA* deletion mutant |
| KpnI-up-comC-F<br>Sal I-up-comC-R | GCA GGTACC GGT ACT TAT CTA GAT ATT GCT AGG GAT<br>GCA GTCGAC ATG ACA ACC TCC TTA TGT AAA TTA TAG T | To amplify upstream of *comC* used for the deletion of *comC* |
| Sma I-down-comC-F<br>Sac I-down-comC-R | GCA CCCGGG GTC ATT GCT GGT TTA GTT GCT TTA A<br>GCA GAGCTC TAC GAA GTC TGC TAA CAA CTC AA | To amplify downstream of *comC* used for the deletion of *comC* |
| Check mutant-comC-F<br>Check mutant-comC-R | CTA GTA TCG GTC TAG ACC ATA CAG ATA<br>TTT GAT GCT GAA GGA TAT CTC AAC C | To verify the *comC* deletion mutant |
| KpnI-up-ssb-F<br>Sal I-up-ssb-R | GCA GGTACC TAA TGC CAG GCG TAT TAA TTA CTG<br>GCA GTCGAC GTA GTG TGT GAT TCA CCT CCT ATG | To amplify upstream of *ssb* used for the deletion of *ssb* |
| Sma I-down-ssb-F<br>Sac I-down-ssb-R | GCA CCCGGG ATC CAA TTA TCC TAA ACA TCC TTA ATA TAC<br>GCA GAGCTC ACT ATT GAT TAG CTA TGC ATA AAT GGC | To amplify downstream of *ssb* used for the deletion of *ssb* |
| Check mutant-ssb-F<br>Check mutant-ssb-R | AGT ATC AGG AAA CGA ACC ATT CTT TCA A<br>GGA ACA TAT CGA GAA TTC CCC GAT ATA T | To verify the *ssb* deletion mutant |
| Gibson-pRIT-gfp-F<br>Gibson-pRIT-gfp-R | GGA TCC GGG AGG CCG TTT<br>CTC TCA GTA CAA TCT GCT CTG ATG CCG | To amplify the linear pRIT used for the reporter plasmid construction by Gibson |
| pRIT-PcomF-F<br>gfp-PcomF-R | ATA TGT GTT CGT GAA TTC CAG TTT GAT CAT AAT TCA GTG TTA CTA TAC ATG GTA CTG<br>AAA CGG CCT CCC GGA TCC TGT TAT TTT ATA TCT TGT TAC ATT ATC CAT TCG ACC CAG TGA TAT | To amplify the promoter of *comF* used for the reporter plasmid constructoin by Gibson |
| pRIT-Pssb-F<br>gfp-Pssb-R | CGG CAT CAG AGC AGA TTG TAC TGA GAG TGC GGT AAT GCC AGG CGT ATT AAT TAC TG<br>AAA CGG CCT CCC GGA TCC TCT CCC GAC AAT TAC GAT TTT ATT TAG CAT GTA GTG TG | To amplify the promoter of *ssb* used for the reporter plasmid construction by Gibson |
| pRIT-PcomC-F | CGG CAT CAG AGC AGA TTG TAC TGA GAG AAG CCT AAC GTT CCA GTG ATA TAT GCT GTT AAA AAT G | |

| | | |
|---|---|---|
| gfp-PcomC-R | AAA CGG CCT CCC GGA TCC TGC AGC TAT AAG ATA ACA ATA CTA CCA AAT GAC AAC CT | To amplify the promoter of *comC* used for the reporter plasmid construction by Gibson |
| pRIT-PcomK1-F | CGG CAT CAG AGC AGA TTG TAC TGA GAG GTA ACG ACA AAT ATA GAA CTT GGG CAT GGA | To amplify the promoter of *comK1* used for the reporter plasmid constructoin by Gibson |
| gfp-PcomK1-R | AAA CGG CCT CCC GGA TCC AAG CAA AAC CTC GCT TTA TTA AGT TTT AAA CC | |
| pRIT-PcomK2-F | CGG CAT CAG AGC AGA TTG TAC TGA GAG GAC CAT CAC GAT ACA TCA TTT CAT TAG TGA | To amplify the promoter of *comK2* used for the reporter plasmid construction by Gibson |
| gfp-PcomK2-R | AAA CGG CCT CCC GGA TCC ATA TAG TAC TCC TCG TAT AAT AAG TTG TTA ATT TAA TGG | |
| pRIT-PsigH-F | CGG CAT CAG AGC AGA TTG TAC TGA GAG AAC AGG TGC AAT TGA ACA TGT ACC AGT TA | To amplify the promoter of *sigH* used for the reporter plasmid construction by Gibson |
| gfp-PsigH-R | AAA CGG CCT CCC GGA TCC GTA TTA AAC TAA CCC CTT CTA TCT AAA ATT TAA GGT TAG | |
| pRIT-X-F | GGC TTA ACT ATG CGG CAT CAG AGC | Used for verifying and sequencing the pRIT constructs |
| pRIT-X-R | GAA TTG TGA GCG GAT AAC AAT TTC ACA CAG G | |
