## Supplemental Figures for "Microaerobic conditions and the complex regulation of natural competence in *Staphylococcus aureus*"

### Supplementary Figures

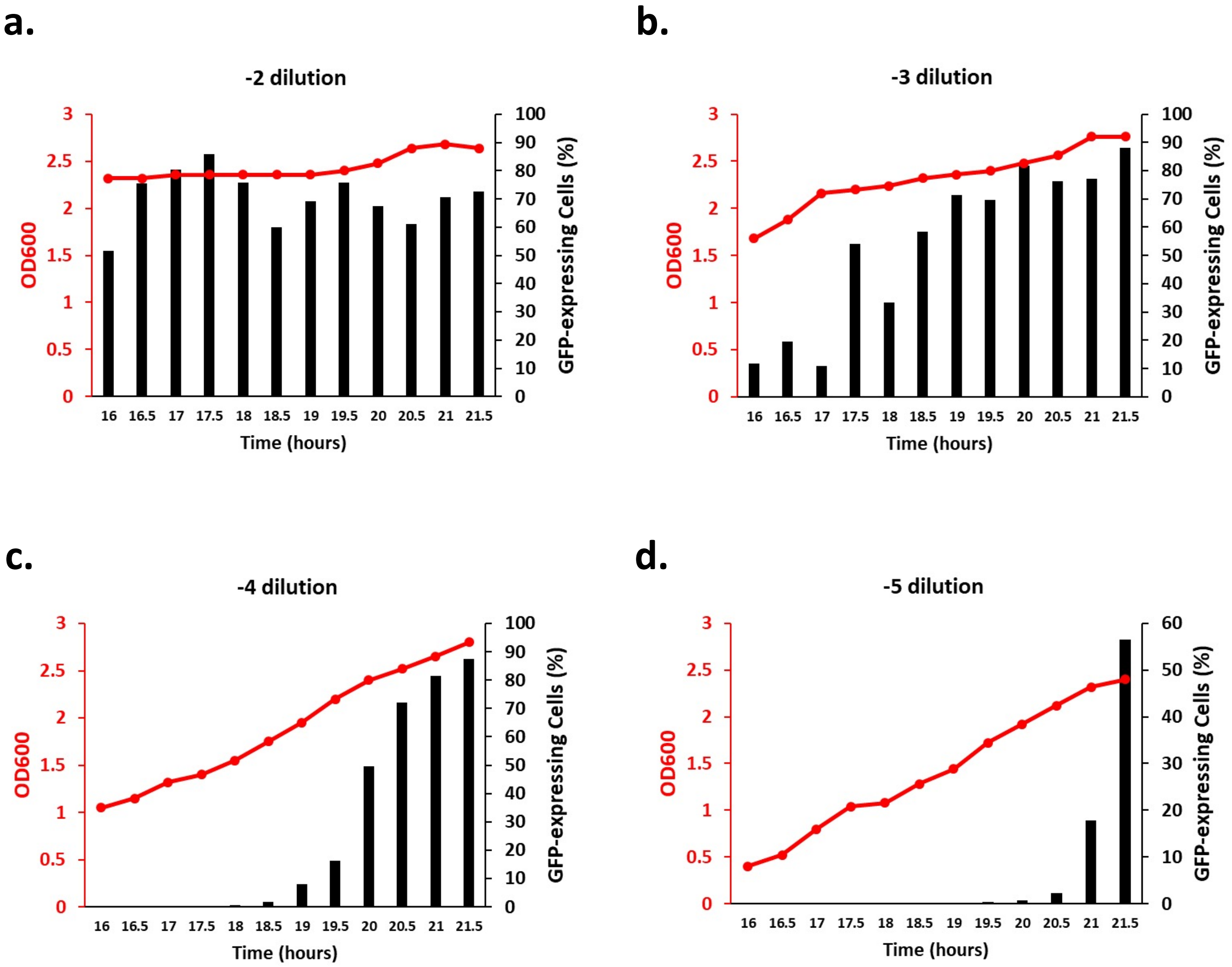


**Supplementary Fig. 1. Development of competence in Individual diluted cultures.**

The growth curve (in red) as well as the evolution of the percentage of competent GFP-expressing cells (black bars) is shown for the -2 (a), -3 (b), -4 (c) and -5 (d) diluted cultures of St29. Following the delay in growth associated to each dilution, the development of competence is also delayed in order to always occur as the cultures approach stationary phase.

a.

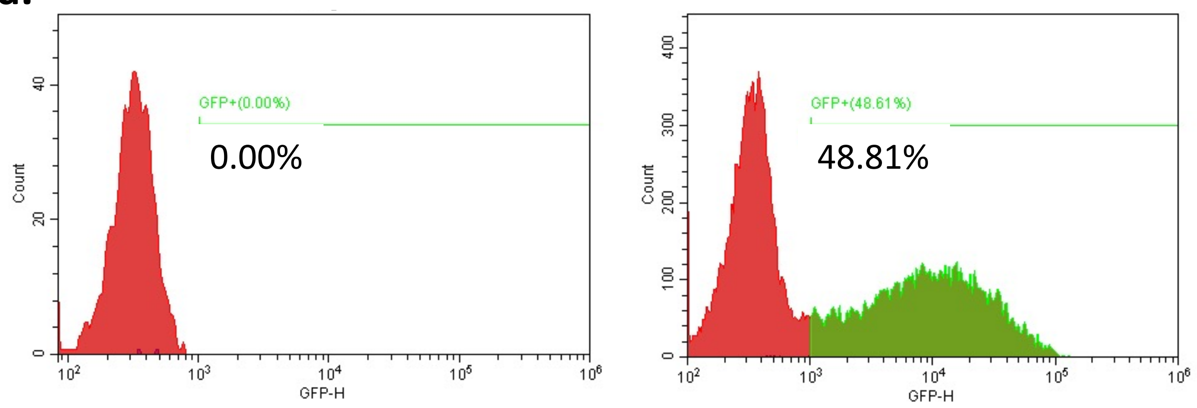


b.

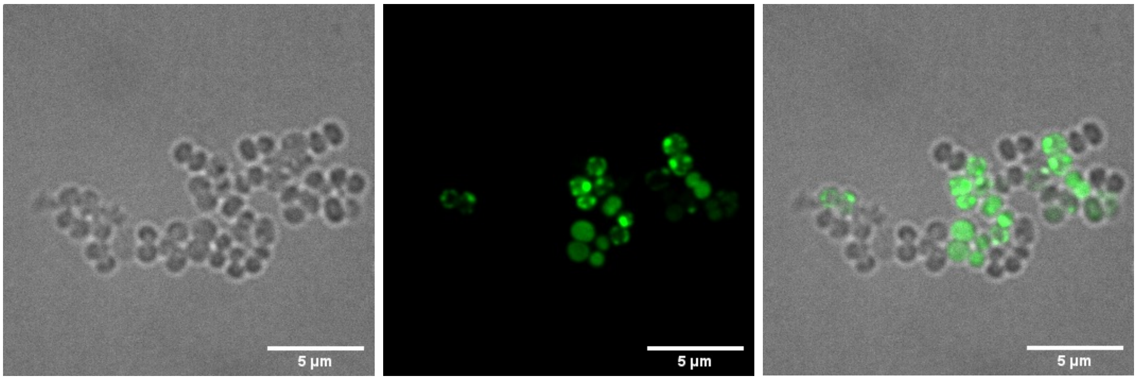


**Supplementary Fig. 2. Flow cytometry and microscopy provide identical percentages of GFP-expressing competent cells (St29)**

a. Classic flow cytometry experiment where the percentage of competent GFP-expressing cells is calculated by comparison of the fluorescence profile of a strain expressing GFP under the control of a competence-induced promoter (in this case, P*comG*, St29) and a strain that does not express GFP (left, St12). The cells showing the maximum auto-fluorescence in the St12 culture, provide the threshold above which cells from the St29 culture are considered as competent (marked in green).

b. Spinning-disk microscopy confirms that the percentage of competent GFP-expressing cells (St29, P*comG-gfp*) is increased with our optimized protocol. In St29, the competent GFP-positive frequency reached 54,1 ± 12,7% (mean ± SD, n=5). Bar = 5 µm.

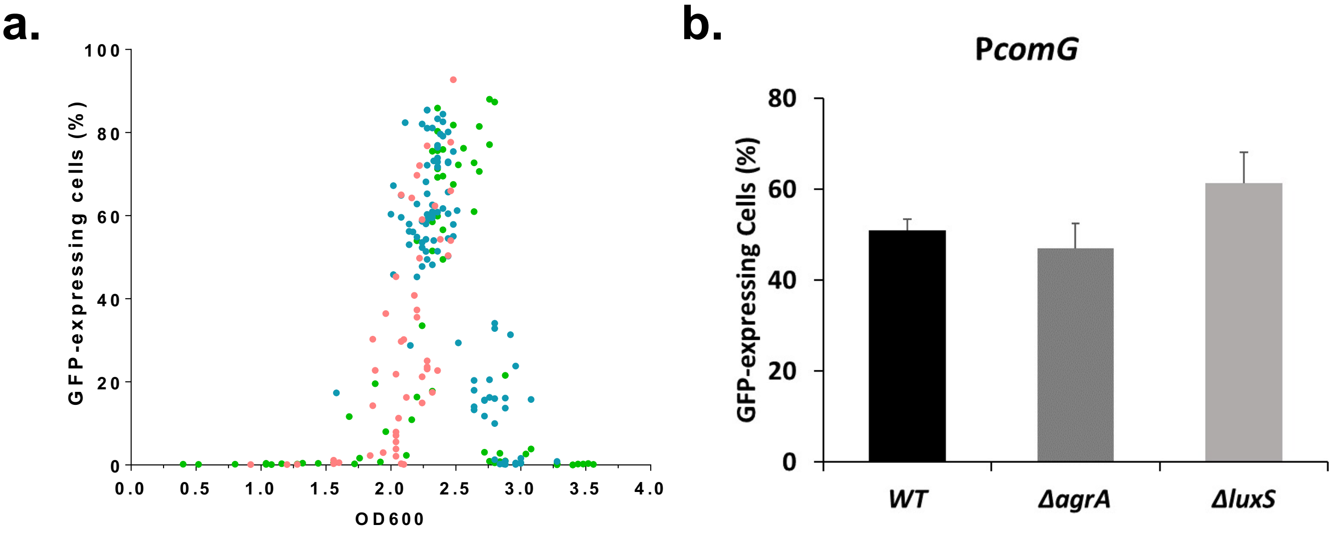


**Supplementary Fig 3. Competence naturally develops at a specific cell density and growth phase in CS2.**

a. Correlation between competence development (percentage of competent cells expressing the GFP under the control of the *comG* promoter, St29) and cell density. The competence window, during which the GFP expression was at its maxima, is allowed when ODs are between 2.2 and 2.6. Data are represented as mean ± SEM of 3 independent experiments (n= 219). Each experiment is materialized by a different color (green, blue or red dots).

b. Percentage of competent cells expressing GFP under the control of the *comG* promoter in wild type (St29) or in the absence of *agrA* (St107) or *luxS* (St123) was determined after 22 hours of growth in CS2. Each experiment in each conditions has been repeated at least 5 times.

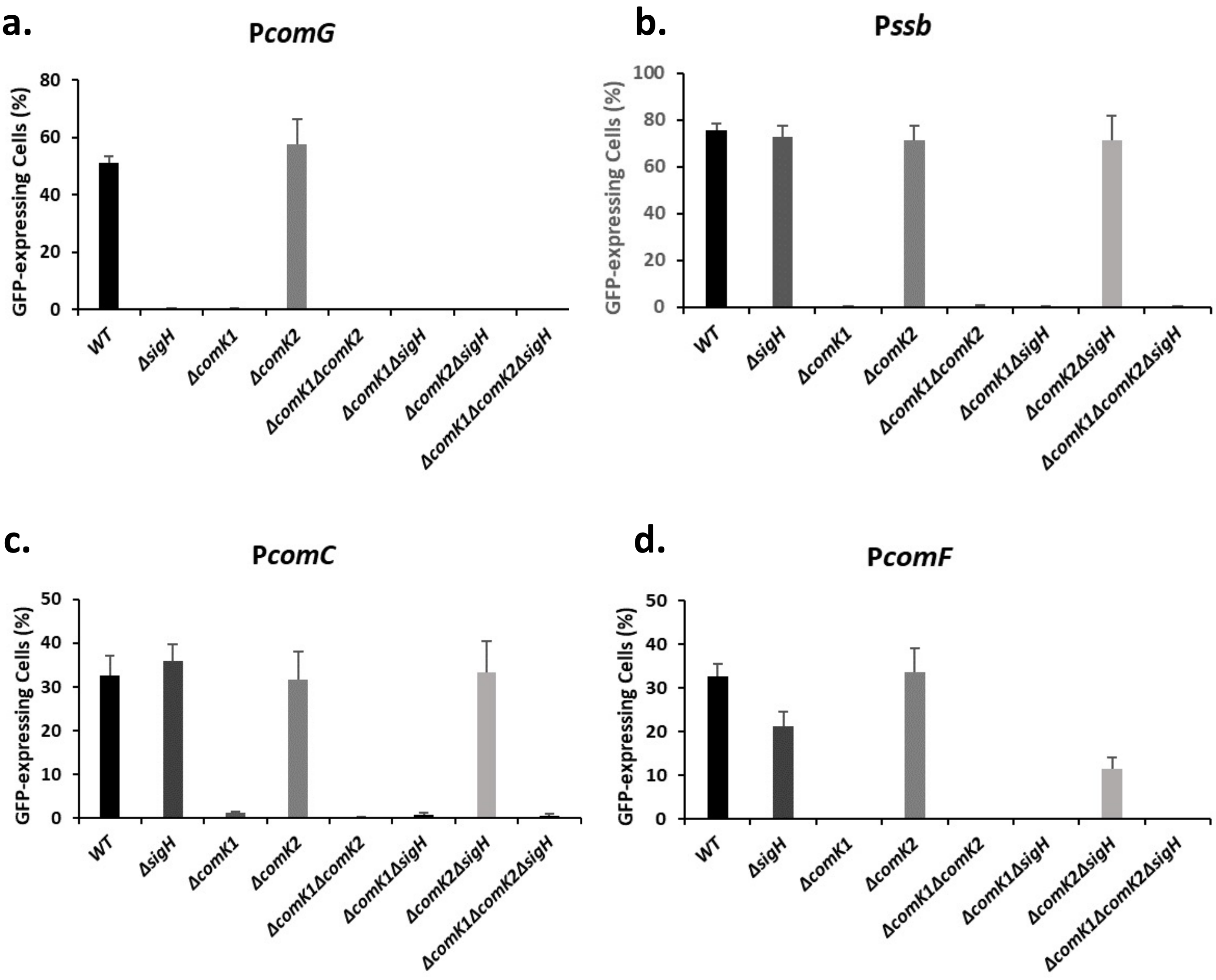


**Supplementary Fig. 4. P*comG*, P*ssb*, P*comC* and P*comF* expression in all mutant strains**

Percentage of competent GFP-expressing cells under the control of P*comG* (St29, St51, St40, St 41, St47, St81, St77 and St69) (a), P*ssb* (St50, St61, St64, St67, St75, St83, St79 and St71) (b), P*comC* (St48, St60, St63, St66, St74, St82, St78 and St70) (c) and P*comF* (St233, St235, St234, St236, St268, St269, St270 and St271) (d) in a wild type background or in the absence of *sigH*, *comK1*, *comK2, comK1/comK2, comK1/sigH, comK2/sigH or comK1,comK2/sigH* was determined after 21 hours of growth in CS2 medium. The results confirm those obtained in Fig. 2. Indeed, independently from the mutant combination considered, as long as one essential regulator is missing, the expression is inhibited.

Each experiment in each condition has been repeated at least 5 times.

| Locus tag N315 | Gene name | Gene function | Δ*sigH* | Δ*comK1* | Δ*comK2* |
|---|---|---|---|---|---|
| SA1899 | *ssb* | single-strand DNA-binding protein | 0.506 | 0.000 | 0.494 |
| SA1374 | *comGA* | traffic ATPase | 0.059 | 0.076 | 0.426 |
| SA1373 | *comGB* | polytopic membrane protein | 0.043 | 0.057 | 0.435 |
| SA1372 | *comGC* | DNA transport machinery, major pilin | 0.051 | 0.051 | 0.520 |
| SA1371 | *comGD* | DNA transport machinery, minor pilin | 0.031 | 0.051 | 0.407 |
| SA1370 | *comGE* | DNA transport machinery, minor pilin | 0.057 | 0.068 | 0.495 |
| SA1369 | *comGF* | DNA transport machinery, minor pilin | 0.030 | 0.059 | 0.404 |
| SA0705 | *comFA* | ATP tanslocase | 0.137 | 0.056 | 0.553 |
| SA0706 | *comFC* | amidophosphoribosyltransferases | 0.101 | 0.043 | 0.536 |
| SA1418 | *comEA* | competence protein, membrane DNA receptor | 0.185 | 0.253 | 0.424 |
| SA1416 | *comEC* | channel protein for DNA binding and uptake | 0.530 | 0.660 | 0.795 |
| SA1486 | *comC* | leader peptidase (prepilin peptidase) | 0.428 | 0.305 | 0.723 |
| SA1128 | *recA* | recombination protein | 0.486 | 0.304 | 0.764 |
| SA1092 | *dprA* | dprA | 0.306 | 0.426 | 0.454 |
| SA1485 | *radC* | DNA repair protein RadC | 0.212 | 0.226 | 0.869 |
| SA0858 | *coiA* | competence protein CoiA | 0.176 | 0.331 | 0.490 |

**Supplementary Fig. 5. Genetic transformation genes expression is induced during natural and controlled by the competence central regulators.**

Overall, we could define three classes of genetic transformation genes: (i) Class I: genes for which SigH and ComK1 are both essential, (ii) Class II: genes (ssb) exclusively controlled by ComK1 (and (iii) Class III genes are controlled by SigH and ComK1 even though their impact is not as important.

However, compared to the GFP reporter strains (**Fig 2a-d**), the impact of the regulators deletion does not always provide the same result by RNA-sequencing. Indeed, the results are confirmed for the *comG, ssb* or *comF* expression, controlled either by SigH and ComK1 or only by ComK1. However, if we consider the *comC* expression, ComK1 and SigH seem to have a small impact through RNA-sequencing **(Table 2)**, while only ComK1 appeared essential using GFP-reporter strains (**Fig 3C**).

Importantly, the differences observed between GFP and RNA-sequencing experiments could be explained by the fact that GFP accumulates during time (because of its stability) while mRNA contents evolve rapidly (because of a high turnover [29]). Therefore, samples of GFP-reporter strains could be taken later into stationary phase, while we extracted the mRNA content at a specific time during the transition to stationary phase. Interestingly, these results suggest that all the genetic transformation genes are not expressed exactly at the same time and that specific regulations (involving one or several regulators) could be at play.

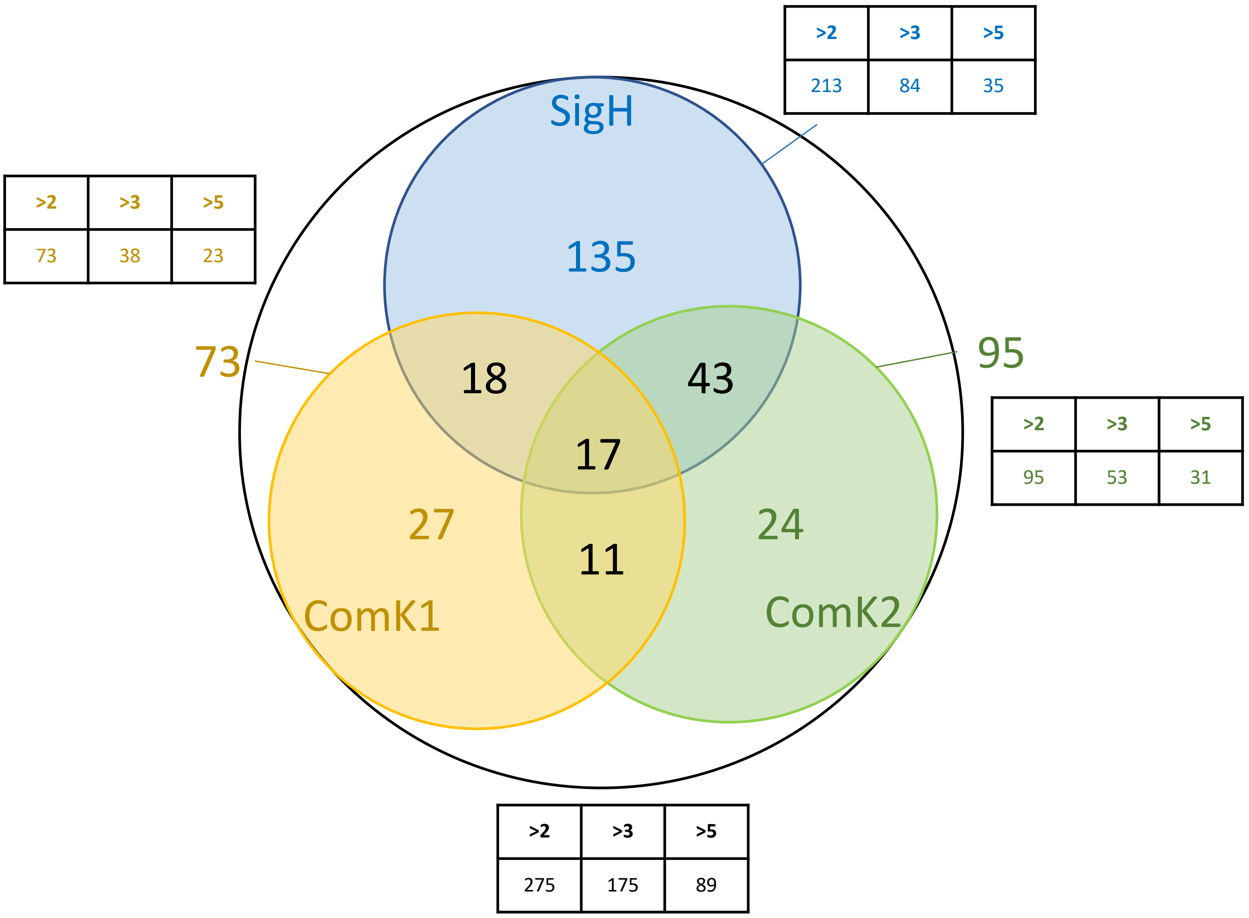


**Supplementary Fig. 6. Global transcriptomic analysis (RNA-sequencing) of competence development in *S. aureus*.**

Each colored circle represent a regulon, for which the expression is increased at least a two-fold factor, and controlled by SigH (in blue), ComK1 (in yellow), by ComK2 (in green) or by the three (outside black circle). The number of genes in each category is shown inside the circles or at the intersection between circles, when the same gene is controlled by more than one regulator. The tables outside show the number of genes for which the expression is increased by a two-fold, three-fold or five-fold factor and controlled by each central regulator (the table color corresponds to the color of the circles) or by the three regulators (in black).

| Gene ID | Function | *Ratio* |
|---|---|---|
| SA2091 | SarY, transcriptional activator | 3,01 |
| purD | Purine metabolism | 3,01 |
| SA0415 | conserved hypothetical protein (ybjB) | 3,02 |
| feoB | major iron permease | 3,02 |
| SA1824 | arsC Arsenate reductase (resistance to arsenic) | 3,06 |
| lytN | cell wall hydrolase | 3,07 |
| SA0632 | Putative lipoprotein | 3,12 |
| SA2150 | Putative hemin transport system permease protein HrtB | 3,14 |
| SA0136 | similar to phosphonates transport permease | 3,14 |
| SA0408 | hypothetical protein | 3,17 |
| clpB | Proteolysis in bacteria, stress | 3,20 |
| SA0704 | hypothetical protein, DegV domain containing prot (operon with comF) | 3,22 |
| SA0530 | alkaline phosphatase, aa (histidine) metabolism | 3,24 |
| SA2090 | fnbB Fibronectin-binding protein, expresion affected by ClpP deletion | 3,24 |
| SA1878 | epiA, lantibiotic epidermin precursor | 3,24 |
| purK | purine biosynthesis pathway | 3,26 |
| SA0333 | Conserved hypothetical protein | 3,30 |
| SA1378 | Conserved hypothetical protein | 3,34 |
| purF | purine biosynthesis pathway | 3,37 |
| SA1822 | Toxin/antitoxin system? | 3,37 |
| SA0838 | glyceraldehyde-3-phosphate dehydrogenase | 3,41 |
| SA0379 | probable transposase | 3,38 |
| modA | Molybdate ABC transporter substrate-binding protein | 3,46 |
| purM | Purine metabolism | 3,47 |
| SA1978 | similar to ferrichrome ABC transporter (permease) | 3,49 |
| SAS028 | sRNA (Bouloc) | 3,51 |
| SA1435 | similar to acetyl-CoA carboxylase accB, fatty acid (lipid) metanolism | 3,55 |
| SA0283 | serine protease? | 3,59 |
| SA0407 | CHP (dipeptidyl aminopeptidases/acylaminoacyl-peptidases) | 3,70 |
| fmhC | secreted bacteriocin | 3,72 |
| SA1636 | pathogenicity island saPI3 | 3,71 |
| butA | Acetoin Reductase | 3,77 |
| radC | DNA repair protein | 3,83 |
| pyrP | uracil permease | 3,90 |
| SA0331 | Efem/EfeO family lipoprotein | 3,96 |
| SA0837 | Amino acid metabolism/transport | 3,99 |
| ndhF | NADH dehydrogenase subunit 5 | 4,14 |
| purN | Purine metabolism | 4,22 |
| SA0412 | hypothetical protein | 4,28 |
| SA0138 | similar to alkylphosphonate ABC tranporter | 4,30 |
| SAS056 | sRNA | 4,34 |
| SA0191 | HlyD family secretion protein | 4,42 |
| SA0413 | hypothetical protein (see SA0412, 0408, 0415) | 4,48 |

**Supplementary Fig. 7. SigH regulon during competence**

List of genes, and their function, for which the expression is induced by more than a three fold factor by SigH. In the last column is provided the exact expression-induction factor associated to each gene under the control of SigH.

The Figure is in two parts (see next figure).

| Gene ID | Function | *Ratio* |
|---|---|---|
| SA1152 | hypothetical protein, hyothetical endotoxin? (see SA1151, 1153) | 4,50 |
| truncated-SA | Lytic regulatory protein truncated with Tn554 | 4,52 |
| SA1418 | comEA, DNA binding and uptake protein | 4,53 |
| SA1249 | hypothetical protein, resistance to vancomycin? | 4,87 |
| purL | Purine metabolism | 4,83 |
| SA0858 | Empbp, secretory extracellular matrix and plasma binding protein, fibrinogen-binding protein (see fnbB) | 4,88 |
| SA2338 | ferrous iron transport protein B homolog (see SA1978) | 5,51 |
| SA1821 | Toxin/antitoxin system? (see Mat&Met of https://europepmc.org/article/pmc/pmc6805077#free-full-text) | 5,53 |
| SA1151 | hypothetical protein | 5,74 |
| lpl4 | lipoprotein | 5,74 |
| SA0705 | comFA; competence protein ComFA | 5,77 |
| uhpT | Sugar phosphate antiporter | 5,95 |
| purQ | Purine metabolism | 6,04 |
| SA2012 | hypothetical protein | 6,38 |
| SA1153 | DNA-dependent DNA polymerase family X | 6,62 |
| valS | valyl-tRNA synthetase, Protein Metabolism biosynthesis | 6,65 |
| SA0919 | purS, phosphoribosylformylglycinamidine synthase, PurS protein | 6,65 |
| SAS040 | Putative uncharacterized protein | 6,96 |
| folC | putative folylpolyglutamate synthase/dihydrofolate synthase | 6,92 |
| purC | Purine metabolism | 7,08 |
| SA0282 | conserved hypothetical protein, Protein secretion system? | 7,36 |
| SA0706 | comFC; competence protein ComFC | 7,38 |
| SA2197 | putative DsbA homologue, protein disulfide bond formation | 7,60 |
| lpl7 | lipoprotein | 8,24 |
| SA0576 | toxin/antitoxin system? | 11,69 |
| SA2198 | putative DsbA homologue, protein disulfide bond formation | 11,80 |
| trpB | tryptophan synthase subunit beta | 12,44 |
| SA1199 | hypothetical protein, similar to anthranilate synthase component I | 12,90 |
| SA1370 | ComGE, Late competence protein ComGE | 13,49 |
| SA1374 | ComGA; competence protein ComGA | 13,53 |
| SA1372 | comGC; competence protein ComGC | 14,69 |
| SA1373 | comGB; competence protein ComGB | 17,92 |
| SA0575 | toxin/antitoxin system? (see SA0576) | 20,20 |
| trpD | Anthranilate phosphoribosyltransferase | 21,41 |
| trpG | anthranilate synthase component II | 22,18 |
| trpC | Indole-3-glycerol phosphate synthase | 22,52 |
| SA1371 | comGD; competence protein ComGD | 23,53 |
| trpF | N-(5'-phosphoribosyl)anthranilate isomerase | 23,80 |
| SA1369 | ComGF, Late competence protein ComGF | 24,56 |
| SA2355 | conserved hypothetical protein, contain Thioredox_DsbH and acetyl-transferase domains | 25,23 |
| SA1979 | similar to toferrichrome ABC transporter | 34,69 |

**Supplementary Fig. 7 (end). SigH regulon during competence**

List of genes, and their function, for which the expression is induced by more than a three fold factor by SigH. In the last column is provided the exact expression-induction factor associated to each gene under the control of SigH.

| Gene ID | Function | *Ratio* |
|---|---|---|
| SA1833 | annotated as a transcriptional activator (Str), Toxin/antitoxin system? | 3,24 |
| SA1486 | ComC, Late competence protein ComC | 3,24 |
| groES | heat shock protein | 3,27 |
| grpE | response to hyperosmotic and heat shock by preventing the aggregation of stress-denatured proteins | 3,28 |
| SA1418 | comEA, DNA binding and uptake protein | 3,30 |
| recA | Recombinase, catalyze the hydrolysis of ATP in the presence of single-stranded DNA | 3,35 |
| xprT | Xanthine phosphoribosyltransferase | 3,48 |
| truncated(radC)-2 | DNA repair protein | 3,59 |
| SA1822 | Toxin/antitoxin system? | 3,78 |
| SA1831 | Toxin/antitoxin system? | 3,82 |
| SA2353 | autolysins | 4,16 |
| SA1978 | similar to ferrichrome ABC transporter | 4,20 |
| lacR | Repressor of the lactose catabolism operon | 4,42 |
| SA1709 | ftnA, Ferritin-like protein | 4,61 |
| SA1820 | operon with SA1821-1822, toxin/antitoxin? | 4,80 |
| hutH | Histidine ammonia-lyase | 5,26 |
| SA1821 | Toxin/antitoxin system? | 5,29 |
| clpB | Proteolysis in bacteria, stress | 6,64 |
| folC | putative folylpolyglutamate synthase/dihydrofolate synthase | 7,23 |
| valS | valyl-tRNA synthetase | 7,53 |
| SA0704 | conserved hypothetical protein, DegV domain containing prot | 8,86 |
| SA2198 | Toxin/antitoxin system? | 9,65 |
| SA1635 | hypothetical protein, Pathogenicity island SaPln3 | 10,28 |
| SA2197 | Toxin/antitoxin system? | 10,16 |
| SA1374 | ComGA; competence protein ComGA | 10,55 |
| SA1370 | ComGE, Late competence protein ComGE | 11,22 |
| SA1369 | ComGF, Late competence protein ComGF | 12,51 |
| SA1373 | comGB; competence protein ComGB | 13,60 |
| SA0705 | comFA; competence protein ComFA | 14,10 |
| SA1372 | comGC; competence protein ComGC | 14,60 |
| SA1371 | comGD; competence protein ComGD | 14,57 |
| SA2355 | conserved hypothetical protein, contain Thioredox_DsbH and acetyl-transferase domains | 15,32 |
| SA0706 | comFC; competence protein ComFC | 17,47 |
| SA1979 | similar toferrichrome ABC transporter | 20,20 |
| SA0575 | Toxin/antitoxin system? | 31,02 |
| SA0576 | Toxin/antitoxin system? | 36,03 |
| SA1898 | hypothetical protein, similar to SceD precursor, able to cleave peptidoglycan and affects clumping and separation of bacterial cells | 721,03 |
| SA1899 | ssb, Single-strand DNA-binding protein | 2533,34 |

**Supplementary Fig. 8. ComK1 regulon during competence**

List of genes, and their function, for which the expression is induced by more than a three fold factor by ComK1. In the last column is provided the exact expression-induction factor associated to each gene under the control of ComK1.

| Gene ID | Function | *Ratio* |
|---|---|---|
| SA1825 | Toxin/antitoxin system? | 3,17 |
| SA0412 | hypothetical protein | 3,30 |
| butA | Acetoin Reductase | 3,42 |
| SA2479 | Polyisoprenoid-binding protein | 3,47 |
| SA1407 | Conserved hypothetical protein, expression affected by ClpP deletion | 3,51 |
| SAS041 | Hypothetical 62 aa protein | 3,53 |
| groES | heat shock protein, 10 kDa chaperonin, CtsR regulon (heat shock) | 3,59 |
| hutH | Histidine ammonia-lyase | 3,63 |
| groEL | heat shock protein, 60 kDa chaperonin, CtsR regulon (heat shock) | 3,67 |
| SA1824 | Toxin/antitoxin system? | 3,78 |
| SA1978 | similar to ferrichrome ABC transporter | 3,88 |
| SA1820 | ribD, Riboflavin biosynthesis protein | 3,89 |
| dnaJ | Chaperone protein DnaJ | 3,90 |
| cysM | cysteine biosynthetic process from serine | 3,91 |
| pyrR | Regulates transcriptional attenuation of the pyrimidine nucleotide | 4,00 |
| SA1823 | Arsenical pump membrane protein | 4,18 |
| SA0916 | purE, phosphoribosylaminoimidazole carboxylase catalytic subunit | 4,22 |
| SA1821 | Toxin/antitoxin system? | 4,27 |
| uhpT | Hexose phosphate transport protein | 4,33 |
| SA1822 | Toxin/antitoxin system? | 4,41 |
| pyrC | Dihydroorotase | 4,84 |
| SA0481 | hypothetical protein, CtsR regulon (heat shock) | 4,91 |
| purK | purine biosynthesis protein | 5,09 |
| purD | purine biosynthesis protein | 5,10 |
| purN | purine biosynthesis protein | 5,29 |
| clpC | CtsR regulon (heat shock) | 5,30 |
| SA0482 | CtsR regulon (heat shock) | 5,33 |
| purH | purine biosynthesis protein | 5,36 |
| dnaK | Chaperone protein DnaK, CtsR regulon (heat shock) | 5,41 |
| purM | purine biosynthesis protein | 5,60 |
| ctsR | regulator of stress response, CtsR regulon (heat shock) | 5,83 |
| purF | purine biosynthesis protein | 5,85 |
| grpE | response to hyperosmotic and heat shock by preventing the aggregation of stress-denatured proteins, CtsR regulon (heat shock) | 5,96 |
| purC | purine biosynthesis protein | 6,00 |
| purL | purine biosynthesis protein | 6,01 |
| SA0919 | Purine metabolism | 6,11 |
| purQ | purine biosynthesis protein | 6,33 |
| SA1596 | aroK, Amino acid biosynthesis | 6,50 |
| hrcA | Heat-inducible transcriptional repressor, CtsR regulon (heat shock) | 6,83 |
| valS | valyl-tRNA synthetase, Protein biosynthesis | 7,54 |
| folC | putative folylpolyglutamate synthase/dihydrofolate synthase | 8,07 |
| trpA | tryptophan biosynthesis | 8,45 |
| SA1199 | hypothetical protein | 8,51 |

**Supplementary Fig. 9. ComK2 regulon during competence**

List of genes, and their function, for which the expression is induced by more than a three fold factor by ComK2. In the last column is provided the exact expression-induction factor associated to each gene under the control of ComK2.

The Figure is in two parts (see next figure).

| Gene ID | Function | *Ratio* |
| --- | --- | --- |
| trpB | tryptophan biosynthesis | 10,37 |
| trpC | tryptophan biosynthesis | 12,91 |
| trpF | tryptophan biosynthesis | 13,70 |
| trpG | tryptophan biosynthesis | 14,94 |
| trpD | tryptophan biosynthesis | 15,68 |
| SA2355 | conserved hypothetical protein, contain Thioredox_DsbH and acetyl-transferase domains | 16,04 |
| clpB | Chaperone protein, CtsR regulon (heat shock) | 19,99 |
| SA1979 | similar toferrichrome ABC transporter | 26,78 |
| SA0576 | Toxin/antitoxin system? | 307,37 |
| SA0575 | Toxin/antitoxin system? | 356,23 |

**Supplementary Fig. 9 (end) ComK2 regulon during competence**
List of genes, and their function, for which the expression is induced by more than a three fold factor by ComK2. In the last column is provided the exact expression-induction factor associated to each gene under the control of ComK2.

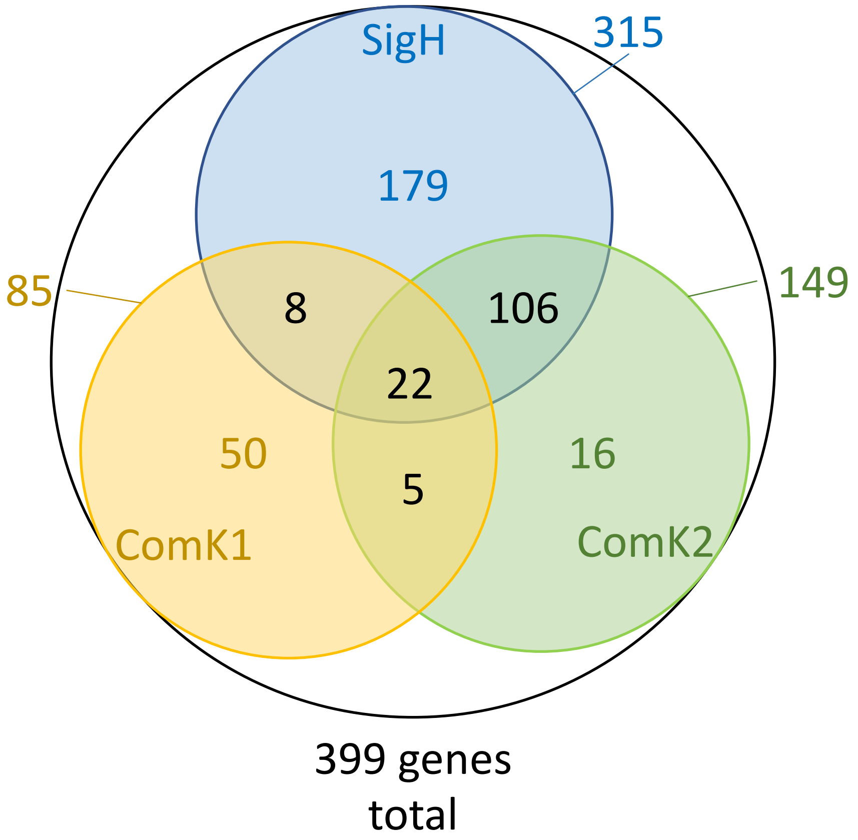


**Supplementary Fig. 10. Genes inhibited during competence**

Each colored circle represent the regulon, for which the expression is decreased at least two-fold, controlled by SigH (in blue), ComK1 (in yellow), by ComK2 (in green) or by the three (outside black circle). The number of genes in each category is shown inside the circle or at the intersection between circles, when the same gene is controlled by more than one regulator. The total number of genes inhibited by SigH, ComK1 and ComK2 during competence development is shown below.

| Gene ID | Function | *SigH* | *K1* | *K2* |
|---|---|---|---|---|
| *capA* | Capsular biosynthesis operon | **4,46** | 2,14 | **4,11** |
| *capB* | Capsular biosynthesis operon | **4,52** | - | 2,71 |
| *capC* | Capsular biosynthesis operon | **5,58** | - | **3,90** |
| *capD* | Capsular biosynthesis operon | **4,22** | - | **3,26** |
| *capE* | Capsular biosynthesis operon | **3,77** | - | **3,45** |
| *capF* | Capsular biosynthesis operon | **3,04** | - | 2,58 |
| *capG* | Capsular biosynthesis operon | 2,84 | - | 2,52 |
| *capH* | Capsular biosynthesis operon | 2,41 | - | - |
| *capI* | Capsular biosynthesis operon | 2,32 | - | 2,09 |
| *capJ* | Capsular biosynthesis operon | 2,41 | - | 2,37 |
| *capK* | Capsular biosynthesis operon | 2,25 | - | - |
| *capL* | Capsular biosynthesis operon | **3,04** | - | 2,31 |
| *capN* | Capsular biosynthesis operon | 2,34 | - | - |
| *capO* | Capsular biosynthesis operon | 2,24 | - | - |
| *clfA* | Clumping factor A | **6,39** | - | **3,31** |
| *clfB* | Clumping factor B | **4,21** | - | **3,46** |
| *coa* | Staphylocoagulase | - | - | 2,49 |
| *fnb* | Fibronectin-binding protein | 2,70 | **3,85** | - |
| *fnbB* | Fibronectin-binding protein | **5,64** | **8,02** | - |
| *geh* | Triacylglycerol lipase | **3,53** | - | - |
| *hlgB* | Toxin (Gamma-hemolysin component B) | - | 2,12 | - |
| *hlgC* | Toxin (Gamma-hemolysin component C) | 2,59 | 2,24 | **3,01** |
| *icaA* | Glycosyltransferase | **3,23** | **3,09** | - |
| *icaB* | Polysaccharide intercellular adhesin deacetylase | 2,14 | 3,15 | - |
| *saeR* | Transcriptional regulator, involved in the regulation of virulence factors | - | - | 2,90 |
| *saeS* | Histidine kinase, involved in the regulation of virulence factors | 2,31 | - | 2,73 |
| *sdrC* | Surface serine-aspartate repeat-containing protein C | 2,24 | - | - |
| *sdrD* | Surface serine-aspartate repeat-containing protein D | **4,15** | - | **3,58** |
| *spa* | Staphylococcal protein A (virulence factor) | **10,28** | 2,11 | **5,38** |
| *splA* | Serine protease | - | - | 2,52 |
| *splB* | Serine protease | - | - | 2,48 |
| *splC* | Serine protease | - | - | 2,27 |
| *splD* | Serine protease | - | - | 2,39 |
| *splF* | Serine protease | - | - | 2,36 |
| *sspA* | Serine protease | 2,39 | - | - |
| *sspB* | Serine protease | 2,03 | - | - |
| *sspC* | Serine protease | 2,32 | - | - |

**Supp. Fig 11. Expression of virulence-related genes is inhibited during competence.**
List of genes involved in virulence, and their function, that were found inhibited during the development of competence. The "inhibition" ratio calculated by comparison of the expression in wild type and mutant strains is presented in the last three columns. Ratios superior to 3 are in bold. 'K1' and 'K2' refer to ComK1 and ComK2.
